# High-resolution vasomotion analysis reveals novel arteriole physiological features and progressive modulation of cerebral vascular networks by stroke

**DOI:** 10.1101/2023.11.05.565680

**Authors:** Yi-Yi Zhang, Jin-Ze Li, Hui-Qi Xie, Yu-Xiao Jin, Wen-Tao Wang, Bing-Rui Zhao, Jie-Min Jia

## Abstract

Spontaneous cerebral vasomotion, characterized by ∼0.1 Hz rhythmic contractility, is crucial for brain homeostasis. However, our understanding of vasomotion is limited due to a lack of high-precision analytical methods to determine single vasomotion events at basal levels. Here, we developed a novel strategy that integrates a baseline smoothing algorithm, allowing precise measurements of vasodynamics and concomitant Ca2+ dynamics in mouse cerebrovasculature imaged by two-photon microscopy. We identified several previously unrecognized vasomotion properties under different physiological and pathological conditions, especially in ischemic stroke, which is a highly harmful brain disease that results from vessel occlusion. First, the dynamic characteristics between SMCs Ca^2+^ and corresponding arteriolar vasomotion are interplayed. Second, compared to previous diameter-based estimations, our radius-based measurements reveal nonisotropic vascular movements, enabling a more precise determination of the latency between smooth muscle cell (SMC) Ca^2+^ activity and vasocontraction. Third, we characterized single vasomotion event kinetics at scales of less than 4 seconds. Finally, following pathological vasoconstrictions induced by ischemic stroke, vasoactive arterioles transitioned to an inert state and persisted despite recanalization. In summary, we developed a highly accurate technique for analyzing spontaneous vasomotion, and we suggest a potential strategy to reduce stroke damage by promoting vasomotion recovery.

## Introduction

Vasomotion has been described as vascular smooth muscle cell (SMC)-initiated spontaneous oscillation that happen relatively independent of pulsatile blood flow^1, 2^. It is characterized by 0.1 Hz sinusoidal changes in vessel diameter^3–5^, regulates cerebral blood flow^6, 7^(CBF).

Spontaneous vasomotion influences both the central nervous system and peripheral systems. In the central nervous system, vasomotion reflecting basal fluctuations in blood oxygenation is related to resting-state neural activity^8, 9^. Recently, vascular optogenetics revealed that enhanced vasomotor activity promotes cerebral spinal fluid influx to the brain parenchyma^10^. As for the peripheral systems, vasomotion modulates fluid filtration pressure in the pulmonary vessels and promotes interstitial fluid clearance in rabbit ear skin^11, 12^. Moreover, the importance of vasomotion for testes showed that vasomotion is essential for normal function^13–15^. These findings highlight the importance of spontaneous vasomotion in tissue homeostasis in both peripheral and central systems.

Vasomotion dysfunction is associated with neuropathies^9, 16^ and oscillatory blood flow at 0.1 Hz has been shown to promote cerebral tissue oxygenation during reduced CBF in human in clinical trials and under high-altitude conditions^17, 18^. Moreover, vasomotion as a driving force can facilitate cerebral paravascular clearance of metabolic waste which is particularly important for elucidating the pathogenesis of Alzheimer’s disease (AD)^8, 9^. These findings highlight that vasomotion is an important new area of research that may support the development of therapeutics for brain diseases such as stroke, AD, and vascular dementia^19^.

However, our current understanding of cerebral vasomotion is not comprehensive. For the generation of vasomotion, the presence of a cellular oscillator is necessary which can normally be visualized under experimental conditions in form of intracellular Ca^2+^ transients^20^. In the central nervous system, SMCs with synchronized Ca^2+^ fluctuations^21^ are responsible for the self-generated movement component of vasomotion, which is independent of external factors such as respiratory rhythms and heart beats^1, 21, 22^. Therefore, it is crucial to explore the correlation and time delay between fluctuation SMC calcium and arteriolar vasomotion for research vasomotion.

The variation of vessel diameter over time is commonly used to characterize vasomotion, while the vessel diameter is defined as distance between bilateral vessel walls and calculated using the half width at half maximum (FWHM) methods^23^. A recently developed vasomotion index integrates multiple features of vasomotion into one general parameter by calculating the area under the diameter-change curve over time; however, this method cannot identify individual events^6^. Moreover, the research on vasomotion focuses on the attributes in the frequency domain due to the complexity of the time domain representation of vasomotion signals^9, 24^. In summary, important aspects of vasomotion remains uncharacterized. A suitable event judgement baseline should be established for basal vasomotion investigations to achieve normalization of vasomotion comparison both in physiological and pathological states, However, neither the baseline F0 minimal nor that F0 average can distinguish vasomotion events under resting-state due to insensitive baseline threshold, although they are frequently used to differentiate large vasomotion signals under a series of stimulation paradigms^25, 26^.

Ischemic stroke is a leading cause of mortality and disability worldwide, characterized by the sudden interruption of blood flow to the brain, resulting in severe neurological deficits. The suppressed vasomotion has been proven to be closely associated with blood flow dysfunction in stroke^27^, and the vasodynamic deficits after successful recanalization have been linked to stroke recurrence and poor recovery after stroke^28^. Mouse models have consistently demonstrated pathological SMC constrictions during occlusion that prevent re-establishment of blood flow^29^. This pathological constriction occurs immediately after the initiation of spreading depolarization (SD) during the hyperacute ischemic phase. However, whether this early SMC constriction leads to secondary, prolonged SMC functional deficits during vasomotion has yet to be determined. Understanding this pathological progression may elucidate stroke characteristics, leading to the development of new methods to prevent delayed neural damage progression.

Here, we integrated multiple algorithms to develop a high-precision analytical method, allowing the quantification of various indices related to vasomotion. These algorithms include vasomotion and calcium kinetics identification through the F0 smoothing algorithm combined with outlier detection, vessel radius definition, time-series curve similarity comparisons through dynamic time warping (DTW), time lag estimation between SMC calcium fluctuations and vasomotion based on sliding cross-correlation values, and arteriolar propagation index quantification through correlation coefficient (CC) maps. This method replicated known vasomotion properties and showed more comprehensive characterization indicators for vasomotion. With this novel method, we identified single vasomotion event kinetics and found a shorter latency than previously known between Ca^2+^ increase and arteriole contraction. Furthermore, we identified the cessation of arteriole vasomotion after stroke. Our results have gained a deeper understanding of vasomotion, and have direct implications for research to cerebral vessel networks and ischemic stroke characteristics.

## Materials and Methods

### Animals

All animal protocols were approved by the Institutional Animal Care and Use Committee (IACUC) at Westlake University. The following mouse strains were used: wild type (C57BL/6J), *SMACreER*, *Ai14*, *Ai47*, *Ai96* (JAX:028866). All mice were bred and maintained in a specific-pathogen-free animal room on a 12-hour light-dark cycle and provided food and water ad libitum at Westlake University Laboratory Animal Resource Center (LARC). All experiments used male and female mice offspring aged 2 to 6 months.

### Experimental timeline

For breading the conditional reporter mice in mural cell, an efficient temporally-controlled transgenic mouse line, the *SMACreER* mice were used^30^. For the tamoxifen-inducible CreER-dependent conditional overexpression systems, we performed tamoxifen (MCE, Cat# HY- 13757A/CS-2870) administration (15 mg/ml dissolved in corn oil, 3 mg daily for three consecutive days through intraperitoneal injection) and induction on adult mice, 3 weeks before in vivo imaging. After cranial window implantation, mice were allowed to recover for 2 weeks prior to imaging, to ensure that surgery-related inflammation had resolved.

### Anesthesia

For surgery and anesthetized two-photon imaging, mice were anesthetized with 1% pentobarbital sodium intraperitoneally (10 ml/kg for mice with a body weight of approximately 25 to 35 g). Core temperature was kept constant at 37°C using a homeothermic heating blanket system during all surgical and experimental procedures.

### Cranial window surgery

A butterfly metal adaptor was glued onto the skull with instance adhesive and dental cement, which was used for head fixation under 2PLSM. The cranial surgery was performed by drilling a 3-mm round window on the anterolateral parietal bone overlying the middle cerebral artery (MCA) territory. Afterward, the cranial window was sealed with a 3-mm round glass coverslip (Warner Instruments, CS-3R, Cat# 64-0720) with the instance adhesive (deli 502 super glue, Cat#No.7146).

### Middle cerebral artery occlusion (MCAO) surgery

Focal cerebral ischemia was induced by the middle cerebral artery occlusion (MCAO) method as described previously^31^. A surgical incision in the neck region was made to expose the right common carotid artery (CCA). After ligating the distal side and the proximal side of the CCA, a small incision was subsequently made between the two ligatures. Then, a silicon rubber-coated monofilament with a rounded tip (Diccol, Cat# 7023910PK5Re) was inserted intraluminally. The monofilament was introduced along the internal carotid artery until the origin of the MCA was occluded. The monofilament was left for 2 hours to prompt transient focal cerebral ischemia. Afterward, reperfusion was performed by withdrawing the monofilament.

### Two-photon in vivo and ex vivo imaging

Mice were live-imaged using a two-photon laser scanning microscope (Olympus, FLUOVIEW, FVMPE-RS) equipped with a cooled high-sensitivity GaAsP PMT detector and an ultrafast IR pulsed laser system (Spectra-Physics, InSight X3, continuously variable wavelength range 680- 1300nm). Pictures were acquired in a 512 × 512 pixels square with a 0.994-µm pixel size under a 25x water-immersion objective (Olympus, XLPLN25XWMP2, NA=1.05). During in vivo imaging measurements (0.625-0.926 Hz), the adult mice were imaged awake during daytime and allowed to move freely on a treadmill while being secured in place by fastening their head in a butterfly metal adaptor. For vascular vasomotion detection in vivo, *SMACreER:Ai14* (tdTomato), *SMACreER:Ai47* (EGFP) mice were used to label the vascular walls. The vascular walls can also be labeled by the fluorescein dye Alexa Fluo633 hydrazide (ThermoFisher, Cat#A30634), FITC-dextran (sigma, Cat#FD2000S, 10mg/ml) or RhoB-dextran (sigma, Cat#R9379, 10mg/ml) through tail intravenous injection^32^. For mural cell cytosolic calcium detection in vivo, *SMACreER:Ai96* mice (GCamp6s reported) were used. A 960 nm wavelength excitation was used navigate SMC, calcium oscillation and vasculature. In MCAO models, the same region was detected before and after ischemic stroke with the frame scan mode under 2PLSM. During ex vivo imaging measurements, the dissected mouse brain was labeled by the Tetramethylrhodamine (TMRM, ThermoFisher Cat#T668) in artificial cerebrospinal fluid (ACSF) at 37℃ for two hours^33^, then the SMC was imaged at 960 nm wavelength under 2PLSM. For relationship research between calcium dynamics and vasomotion, the built-in line-scan mode of the two-photon microscope was used to take time-lapse information (100-200 Hz).

### Vessel diameter and corresponding SMC Ca^2+^ signal measurement

All time-lapse pictures were analyzed in Fiji (version 2.3.0/1.53f) and MATLAB (version R2021a; MathWorks) using custom-written scripts. The vessel-containing images were loaded into Matlab and then underwent gaussian filtering to reduce spatial noise. The diameter of the vessel was determined as FWHM of the reslice profile^23^, and the diameter change was obtained by sorting the diameter obtained by each time stack. In the calcium-signal-containing images, regions of interest (ROIs) were drawn on several SMCs according to their outline, and the change in calcium signal under time series was reflected by changing of the average green fluorescence brightness of each SMC.

### Vessel radius definition

For radius calculation, the midpoint line was generated by dividing individual diameter through two along the time series, based on the calculated FWHM. To reduce the influence of errors in the data and extract a robust and appropriate radius boundary, the centerline is fitted by midpoint line using the least-square method (LSM) method^34^. The centerline was defined as one boundary of the side1 radius and side2 radius and the unilateral FWHM as the other boundary for radius calculation, and eventually the radius was subsequently calculated.

### Power spectrum analysis

The Fast Fourier Transform (FFT) is employed to calculate the power spectral density of vessel diameter and radius, and mural cell calcium data. The mice vasomotion and calcium signals (FFT length: 200, the overlap: 50%) under awake/anesthesia and the filtered power of the ultra-low frequency spectrum (0-0.3 Hz) were shown.

### Pearson correlation coefficient and propagation index analysis

Previously, Pearson correlation coefficient (CC) and propagation analysis was implemented not only to identify the functional connectivity between ultra-slow single-vessel bold spatiotemporal dynamics and neuronal intracellular calcium signals^24^, but also to search spontaneous vasomotion propagates along pial arterioles in awake mouse brain^26^. By choosing intervals of 2.5 μm, the vasomotion is calculated at each position ±30 μm from the core position along the arteriole. Compare the CC between each position vasomotion and the core position vasomotion, the corresponding curve can be obtained. The average CC maps is shown as CC comparison of each vasomotion statistic position. The vasomotion synchronization 3D model, simultaneously presents the propagation parameters of time, distance, filtered amplitude ratio, and CC curve of vasomotion.

### Time lag estimation between SMC calcium fluctuation and vasomotion

Based on the results of SMC calcium dynamic and arteriolar vasomotion from time-lapse images, there is a negative cross-correlation between the two^6, 21^. The time lag between SMC calcium and arteriolar vasomotion was presented as the moment when cross-correlation curve arrived the minimum correlation.

### Dynamic time warping analysis

Dynamic time warping (DTW) is widely used as a similarity measurement to find an optimal alignment two given (time-dependent) sequences under certain restrictions^35^. Because of the universal adaptation and robust advantages of DTW in similarity comparison, this method can be combined with CC to describe the similarity of different vasomotion motion patterns. Here, based on the conclusion that both internal and external radius can be effectively utilized to investigate the properties of spontaneous vasomotion, the control group which means the same vasomotion motion patterns was defined as the paired DTW distance comparison between time series of internal and external diameter. Then the DTW distance between time series of side1 radius and side2 radius was calculated which had significant difference compared to the control group to give vital quantitative evidence that asymmetrical motion between the two sides of arteriolar vasomotion.

### Calcium and vasomotion index calculation

For myogenic spontaneous vasomotion characterization, all data used for extracting the vasomotion index must follow a normal distribution. The vasomotion index was defined as quantitative indicators on the total number of vasomotion events^36^. Here, we defined that the eighth smallest number in a forty sliding window producing a variable F0 time series, referred to as the “F0 smooth”. When the amplitude of vasomotion exceeded the double SD of baseline, it is defined as a single event. The kinetic quantification index of events including the calcium index, diameter index, and radius index. These indexes all contained parameters of frequency, SD of peak intervals (interval SD), absolute amplitude and amplitude ratio.

### Rank distribution of blood vessels

During analysis of surface arterioles, the three breakpoints were employed to categorize the vessels into four diameter ranks with locations at 23.05 μm, 32.18 μm and 38.83 μm. These numbers corresponded to the quartiles value in normalized histograms for arterioles segments, were the natural point to divide the data^37^.

## Quantification and statistical analysis

The numerical data concealed in raw digital images were extracted and run on the software of Fiji or MATLAB. All statistical analyses and graphical illustrations were performed using GraphPad Prism 8 software (version 8.3.1, California, USA). All statistical tests were two-tailed, and significance was assigned at p < 0.05. Normality between group samples were assessed using the Kolmogrov-Smirnov test, Lilliefors test, Jarque-Bera test and Chi square Goodness-of-Fit test. When normality between sample groups was achieved, a paired Student’s t-test were used. The Pearson’s correlation values would only be recorded in the statistical data if the p<0.05. The data with error bars are displayed as the means ± SEM. No blinding and randomization design was needed in this work.

## Code availability

All image processing and analysis were performed in MATLAB. The codes consist of seven parts: Diameter radius detection, Frequency calculation, Propagation index calculation, 3D synchronization analysis, Time lag calculation, Dynamic time warping analysis and vasomotion index calculation, which are available at https://github.com/JialabEleven/Vasomotion_analysis.

The detailed explanation of algorithm-related formulas can be found in supplementary data.

## Results

### A high-precision analytical method for vasomotion and achieved parameters

In this study, we focused on the pial arteriolar networks, which are known to exhibit distinct vasomotor activity. The identification of these networks was facilitated by the distinctive morphological characteristics of their mural cells, which were demonstrated in Rosa26-reporter mice crossed with *SMACreER* mice. Specifically, the tamoxifen-inducible expression of Cre under control of the SMA promoter allowed for the labeling of nearly all arterioles and a very small proportion of large venules ( > 100 µm, Supplemental Fig. 1). Intriguingly, our observations consistently revealed that arterioles were positioned above venules when they intersected spatially. This phenomenon was also observed in *SMACreER:Rosa26*-reporter mice (Supplementary Fig. 1A, Movie 1). Consequently, this characteristic allowed us to reliably distinguish pial arterioles from venules in wild-type mice that received intraluminal dye injections.

To perform a comprehensive description of vasomotion, we performed an eight-step process to analyze the time-lapse images and assessed nine parameters related to spontaneous vasomotion, as specified below. In step 1, we generated a time-stack kymograph for each vascular segment utilizing the reslice function in ImageJ with linear regions of interest (ROIs) that were strictly perpendicular to the longitudinal axis of the targeted vessel. Only well-focused vessel segments were chosen for vasomotion analysis, accounting for the curvature of pial vessels and excluding all out-of-focus vessels (Supplementary Fig. 2). This criterion is important for accurate vasomotion studies.

In step 2, we digitized the brightness of each pixel along each horizontal line in the kymograph to form a time stack matrix of the image, and the image stacks were processed in MATLAB with Gaussian filtering to reduce spatial noise (Fig. 1). In step 3, we determined the FWHM coordinates of both boundaries to calculate the vessel diameter for each image stack and digitally assessed dynamic variations in vessel diameter over time (step 4).

**Figure. 1.**
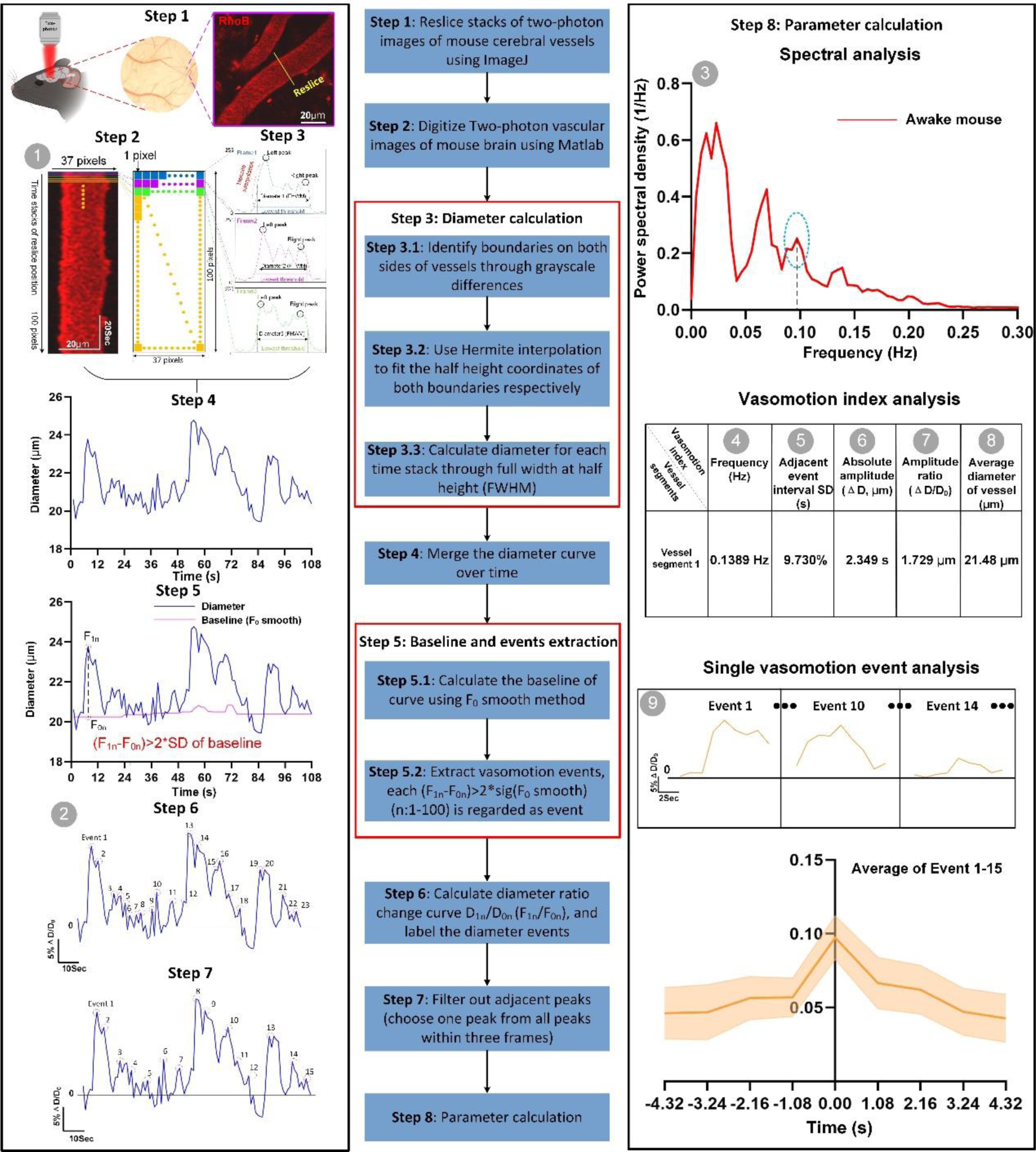
Flow chart of developing a comprehensive index for quantitatively and precisely characterizing spontaneous vasomotion. In vessel-containing images, time stacks of line ROIs are drawn on several vessel segments (The yellow solid line represents the reslice position, on average 2-8 segments per FOV), where pixel brightness along a line orthogonal to the length of the blood vessel was extracted. Image stacks in Tiff format are loaded into Matlab and total nine vasomotion parameters are output (The grey number ball indicates the detailed information of each parameter). These parameters are numbered in numerical order, representing as kymograph (1), time series diameter change curve (2), spectral analysis of diameter change curve (3), frequency of vasomotion (4), the standard deviation (SD) of interval times between two adjacent events (5), the absolute difference in vascular diameter change (6), the amplitude of difference in vascular diameter change (7), the amplitude ratio of vasomotion (8), the average diameter of vessel of time series and analysis of single events (9), respectively. All the results are exported into Excel.

In steps 5 and 6, the baseline diameter (D0) variation over time was calculated using the D0 smoothing method. Initially, we adopted three methods that are commonly used to determine the baselines of the time series of the Ca^2+^ fluorescence (F) intensity fluctuations in neurons under stimulated conditions, namely, F0 average, F0 minimal, and F0 smooth^36^. The last method requires a time-rolling algorithm, while the first two methods simply use the average change or the smallest change as a baseline. Under basal conditions with no external stimulation, vasomotion events are difficult to differentiate with two methods that generate a fixed baseline (D0 minimal and average) when applying usual criterion for identifying a signal, namely, that the signal should exceed double SD of the diameter curve (Supplementary Fig. 3). In contrast, we detected more peaks when we employed the F0 smoothing method. Additionally, to minimize the number of false positives (step 7), we filtered the above results by selecting one event when two events were detected within three consecutive frames. To our knowledge, we are the first to utilize the time-rolling strategy in basal vasomotion analysis.

In addition, we applied traditional frequency and time domain analysis approaches to quantify nine parameters of spontaneous vasomotion events (Fig. 1), including kymographs, time series of the diameter ratio change, and spectral analyses of the diameter change curve. Vasomotion can be divided into five components: the frequency of vasomotion, the standard deviation of the time interval between two adjacent events, the absolute difference in the vascular diameter change (ΔD), the amplitude ratio of the vasomotion (ΔD/D0), and the average diameter of vessel. Importantly, a novel contribution by elucidating the vasomotion dynamics, introducing the concept of single vasomotion cycles. In summary, we developed a procedure to comprehensively quantify spontaneous cerebral vasomotion events with high precision.

### Freshly dissected mouse brains exhibit altered but noticeable arteriolar spontaneous vasomotion

To study vasomotion dynamics of arterioles, we compare arteriolar vasomotion index between freshly dissected mouse brains and awake mouse brain. Although isolated arterioles contract in vitro due to SMC-generated myogenic forces^38^, two key uncertainties remain in ex vivo brain. Firstly, it is unclear whether the ex vivo vasomotion resembles the ∼ 0.1-Hz vasomotion in awake brains. Secondly, the quantitative differences of ex vivo vasomotion compared to in vivo vasomotion are still unknown. We obtained vasodynamics data through two-photon microscopy, examining both freshly dissected mouse brains and head-fixed awake mice positioned on a treadmill with a surgically opened cranial window (Fig. 2A). Ultimately, the effectiveness of vasomotion properties methods (Fig. 1) will be assessed by comparing these two conditions.

**Figure. 2.**
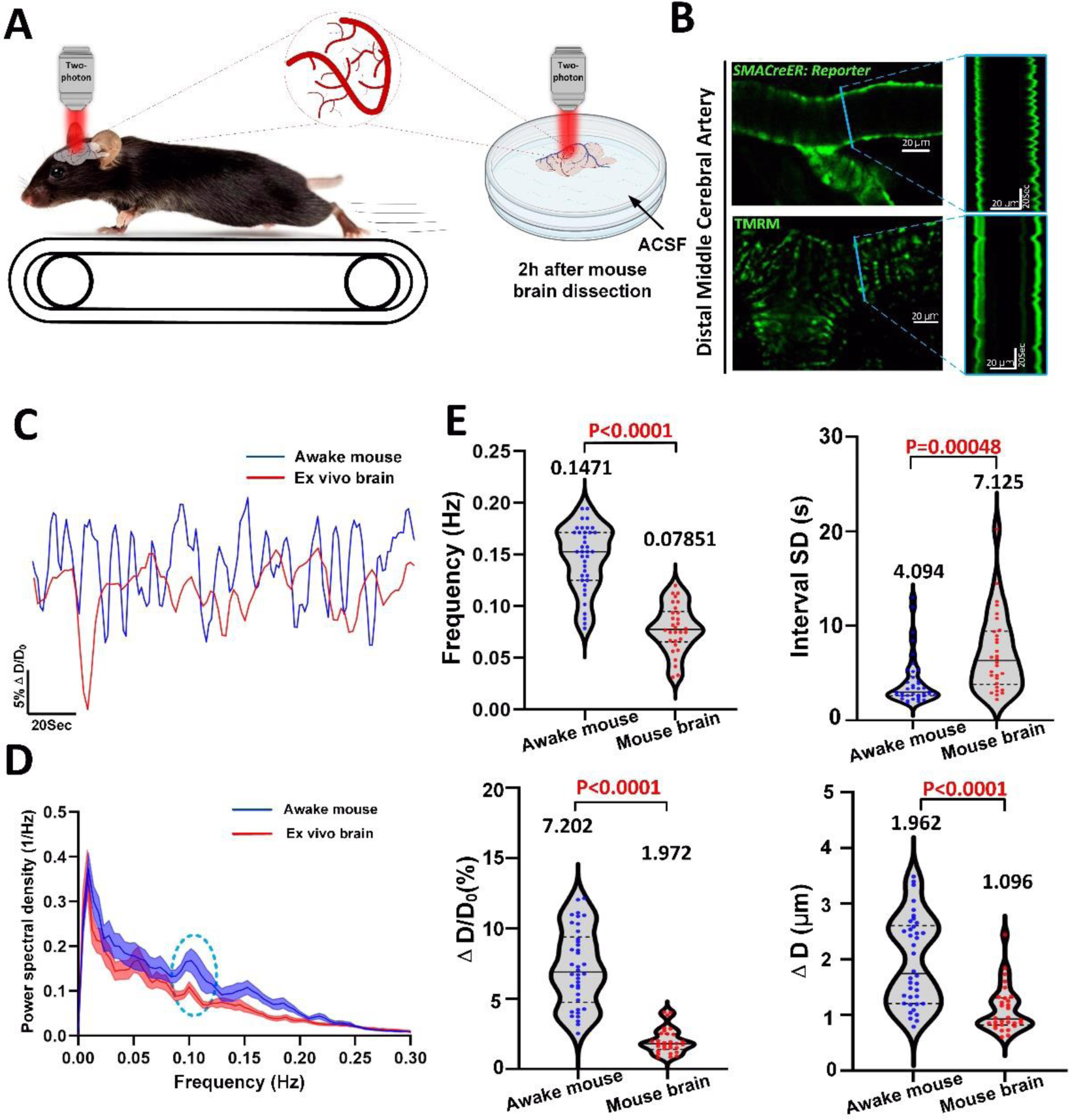
Vasomotion index analyses reveal freshly dissected mouse brains exhibiting altered but noticeable arteriolar spontaneous vasomotion compared to awake mouse brains. (A) Experimental design of the spontaneous vasomotion detection in awake mouse and ex vivo mouse brain. (B) Representative still-frame in vivo two-photon (2P) microscopy images and kymographs of cerebral arteriole through a cranial window in an awake head-fixed mouse and ex vivo mouse brain. The blue solid line represents the resliced position. (C) Representative time-series amplitude ratio (ΔD/D0) changes trace of awake mouse arteriole (blue solid line) and ex vivo mouse brain arteriole (red solid line). (D) The averaged Fourier plot across awake mouse cerebral diameter change and ex vivo mouse brain cerebral diameter change (n=15 arterioles segments in 4 awake mice) revealed a broad range of ultra-low frequencies (<0.3Hz), with a distinct peak centered at around 0.1 Hz. Shaded areas represent SEM. (E) Vasomotion index analysis between awake mouse (n=38 arterioles segments in 4 awake mice) and ex vivo mouse brain (n=30 arterioles segments in 4 mouse brains). Top left represents frequency of vasomotion, top right represents the SD of interval times between two adjacent events (interval SD), bottom left represents the amplitude ratio Δ D/D0 of vasomotion and bottom right represents the absolute difference in vascular diameter change ΔD.

The kymographs generated in blue lines revealed differences in vasomotion profile under these two conditions (Fig. 2B, Movie 2-3). The time series of ΔD/D0 further corroborated this disparity (Fig. 2C). Although different in these aspects, Fourier analysis revealed that dissected brains kept in ACSF for two hours without an external supply of oxygen resembled characteristic peaks around 0.1 Hz observed in vivo (Fig. 2D). The vasomotion index analyses indicated that all four aspects of vasodynamics robustly decreased in ex vivo brains (Fig. 2E, Table 1). Altogether, these data not only demonstrate that the local control of SMCs over arteriole diameter can be up to two hours ex vivo but also support that vasomotion analyses sufficiently precise to discern differential vasomotion characteristics under different conditions.

### Vasodynamic changes derived from both internal and external diameters can be used to characterize spontaneous vasomotion

Depending on the specific experimental conditions, some previous reports have focused on internal diameter^9, 37, 39^, while others have measured external diameter^6, 40^. Consequently, we considered whether the vasomotion features extracted from these two diameters are comparable. All figures utilize green color to represent external diameter and red color to represent measured internal diameter of the lumen space within the vessels (Fig. 3A-3C).

**Figure. 3.**
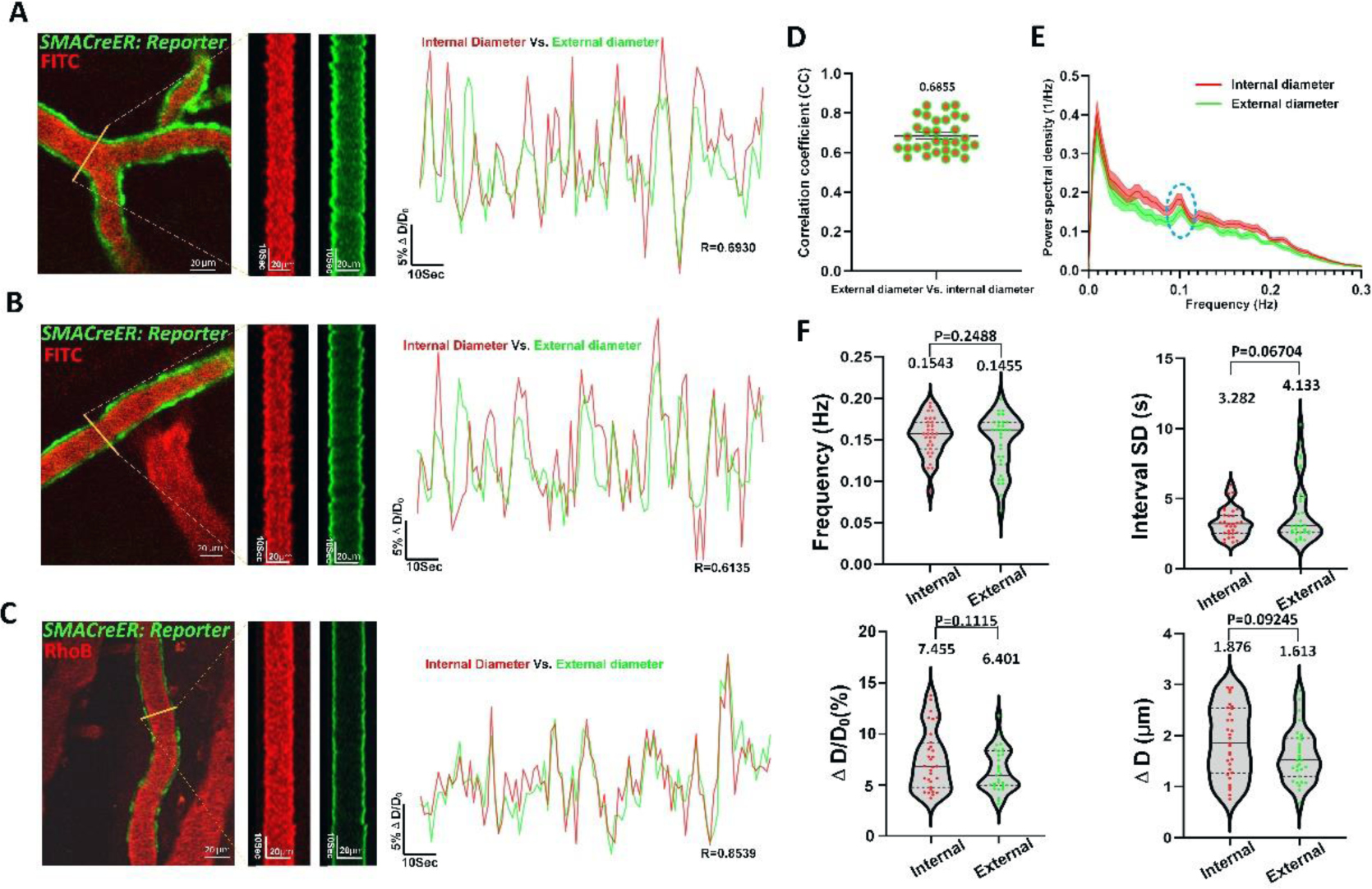
All dynamic changes in internal and external vascular diameter can be used to characterize spontaneous vasomotion. (A)-(C) Representative three levels still-frame images and kymographs of the same cerebral arteriole internal (red) and external (green) diameter and corresponding medium, low and high correlation coefficient (CC) of time-series external-internal diameter amplitude ratio ΔD/D0 changes curves, respectively. The yellow solid line represents the reslice position. (D) Quantification of CC values between same arterioles segments internal and external diameter vasomotion (n=31 arterioles segments in 8 awake mice). (E) The averaged Fourier plot across awake mouse cerebral internal and external diameter change of same arterioles (n=31 arterioles segments in 8 awake mice) revealed a broad range of ultra-low frequencies (<0.3Hz), with a distinct peak centered at around 0.1 Hz. Shaded areas represent SEM. (F) Vasomotion index analysis of cerebral arteriole between internal and external diameter (n=31 arterioles segments in 8 awake mice). Top left represents the index of frequency, top right represents the SD of the adjacent vasomotion event interval, bottom left represents the index of amplitude ratio Δ D/D0 and bottom right represents index of absolute amplitude ΔD. The reporter mice used in our study included Ai14, Ai47, and Ai96 mice which external diameter are uniformly labeled as green color and internal diameter are labeled as red color.

In total, we examined 31 arteriole segments from eight mice to address this issue. Spontaneous contractions and dilations of arterioles were observed in both external and internal diameter kymographs (Fig. 3A-3C). Moreover, the time series of ΔD/D0 revealed high correlations between these two types of diameters (Fig. 3D), and a 0.1-Hz peak was observed in the spectral analysis (Fig. 3E). Moreover, our findings showed no discernible differences in the vasomotion index (Fig. 3F, Table 2). Based on these comparisons, we conclude that internal and external diameters can both be effectively utilized to investigate the properties of spontaneous vasomotion (Movie 4). This is particularly important when there is a need to retrospectively compare features between data that have already been obtained solely from the external or internal diameter.

### The radius vasomotion index reveals that the circular arteriolar wall does not isotropic contract and dilate in an isotropic manner

In the kymograph and time series data analyses performed in this study as well as previously published representative kymographs of vasodynamics obtained under various conditions, including basal, neural, and chemical stimuli conditions^6^, we observed that the two sides of the kymograph tended to be asymmetric (Fig. 2B). Therefore, we developed a radius index to characterize this nonisotropic contractility of arterioles.

Next, we verified that the internal and external radii, as well as the diameter, can be employed to investigate spontaneous vasomotion properties (Supplementary Fig. 4, Table 2). To accomplish this, we compared the time series of each radius to that of the diameter and observed mismatched dilations and contractions (Fig. 4A). In contrast, mismatches between the external and internal radii were seldom observed (Fig. 4B, Movie 5).

**Figure. 4.**
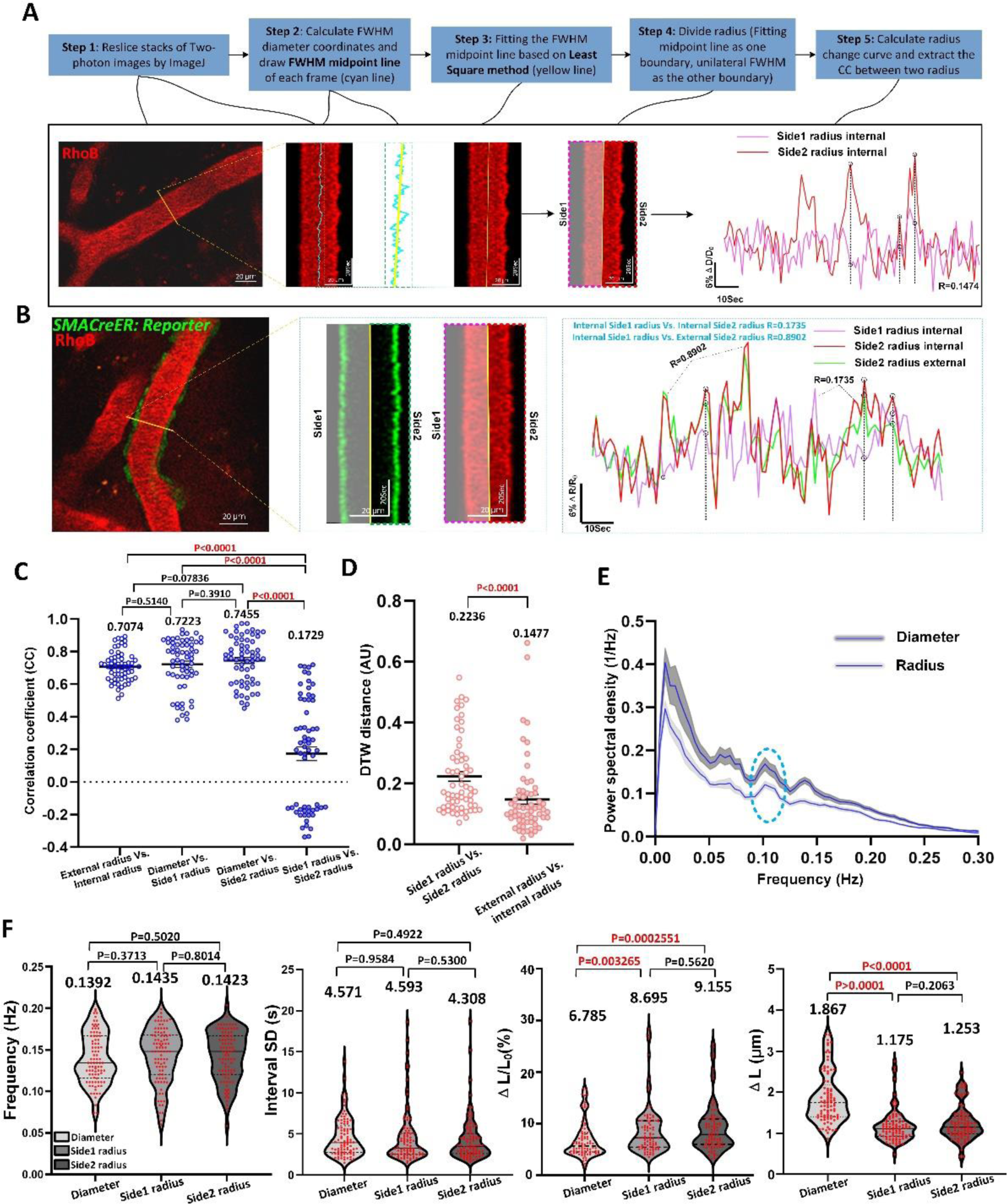
Radius vasomotion index reveals an asymmetrical movement between the two sides of the arteriolar wall at the maximum diameter section. (A) Flow chart of radius boundary definition. The yellow solid line represents the reslice position and the black dotted line represents the mismatched vasomotion. (B) Representative still-frame image and kymograph of cerebral arteriole internal (red) and external (green) radius. Right shows the CC values comparison of side1 radius and side2 radius amplitude ratio with internal and external radius amplitude ratio as reference. Shaded areas represent radius of the other side. (C) The statistic graph of CC value comparison reveals the correlation between internal and external radius amplitude ratio (n=62 arterioles segments in 8 awake mice), diameter and side1 radius (n=62 arterioles segments in 9 awake mice), diameter and side2 radius (n=62 arterioles segments in 9 awake mice), side1 radius and side2 radius (n=62 arterioles segments in 9 awake mice). (D) Dynamic time warping (DTW) comparison reveals the difference of time-series radius change ratio curves between side1 and side2 radius (n=62 arterioles segments in 9 awake mice) of the same cerebral arteriole with internal and external radius (n=62 arterioles segments in 8 awake mice) as reference. (E) The averaged Fourier plot comparison across diameter (n=31 arterioles segments in 9 awake mice) and radius (n=62 arterioles segments in 9 awake mice) in awake mouse revealed a broad range of ultra-low frequencies (<0.3Hz), with a similar distinct peak centered at around 0.1 Hz (Blue round solid line). Shaded areas represent SEM. (F) Vasomotion index analysis between diameter and radius (n=89 arterioles segments in 9 awake mice).

Two types of disparities between the time-series curves should be considered: temporal disparities and amplitude differences. Thus, we examined the mismatches between the curves by combining the dynamic time warping (DTW) method with correlation coefficient (CC) analysis to gain a more comprehensive understanding of arteriolar wall movement. A higher DTW score signifies a greater distance (and thus less similarity) between two analyzed objects (Supplementary Fig. 5).

There are four sets of CC values of different length amplitude ratio were compared separately (Fig. 4C, Table 3), The CC obtained from external radius highly positively correlates with its corresponding internal radius. Similarly, the CC values of diameter were highly positively correlates with its corresponding radius. In contrast, the side1 radius did not correlate with side 2 radius (0.1729±0.04281, p<0.05). Moreover, the DTW score obtained by comparing the two radii corresponding to a single diameter was significantly higher than that obtained by comparing the side 1 radius with its corresponding external radius (Fig. 4D and Supplementary Fig. 5). This finding supports the asymmetry between the opposite arteriole walls.

Despite this asymmetry, the radius analysis showed a peak at approximately 0.1 Hz, like the results of the diameter analysis (Fig. 4E). Additionally, the radius index was generally comparable to the diameter index, except for two parameters: percent of length changes (ΔL/L0) and absolute difference in the length changes (ΔL) (Fig. 4F, Table 4). This difference in length can be attributed to the fact that the radius is half of the diameter. Overall, these data highlight that the radius-based analysis has unique advantages over diameter-based analyses for study aims, such as the extraction of time delay between fluctuation SMC calcium and arteriolar vasomotion, which will be discussed in detail in the upcoming sections.

### Arterioles exhibit suppressed spontaneous vasomotion and weakened vasomotion propagation index under anesthesia

To examine whether cerebral arterioles exhibit prominent 0.1-Hz vasomotion before and after anesthesia, we performed a paired comparison of the same arterioles in mice in awake and anesthetized states; the latter state was induced by intraperitoneal administration of pentobarbital sodium (10 ml/kg). It is known that anesthesia does not deplete arteriole spontaneous vasomotion but dilates the resting diameter^21, 41^, but it is still unclear to what extent anesthesia can induce. Furthermore, the differential vasomotor responses of arterioles to anesthesia have not been quantitatively characterized.

The results showed that 25 arteriole segments from four mice with diameters ranging from 21.87 µm to 74.46 µm were all dilated when the mice were deeply anesthetized (Fig. 5A-D).

**Figure. 5.**
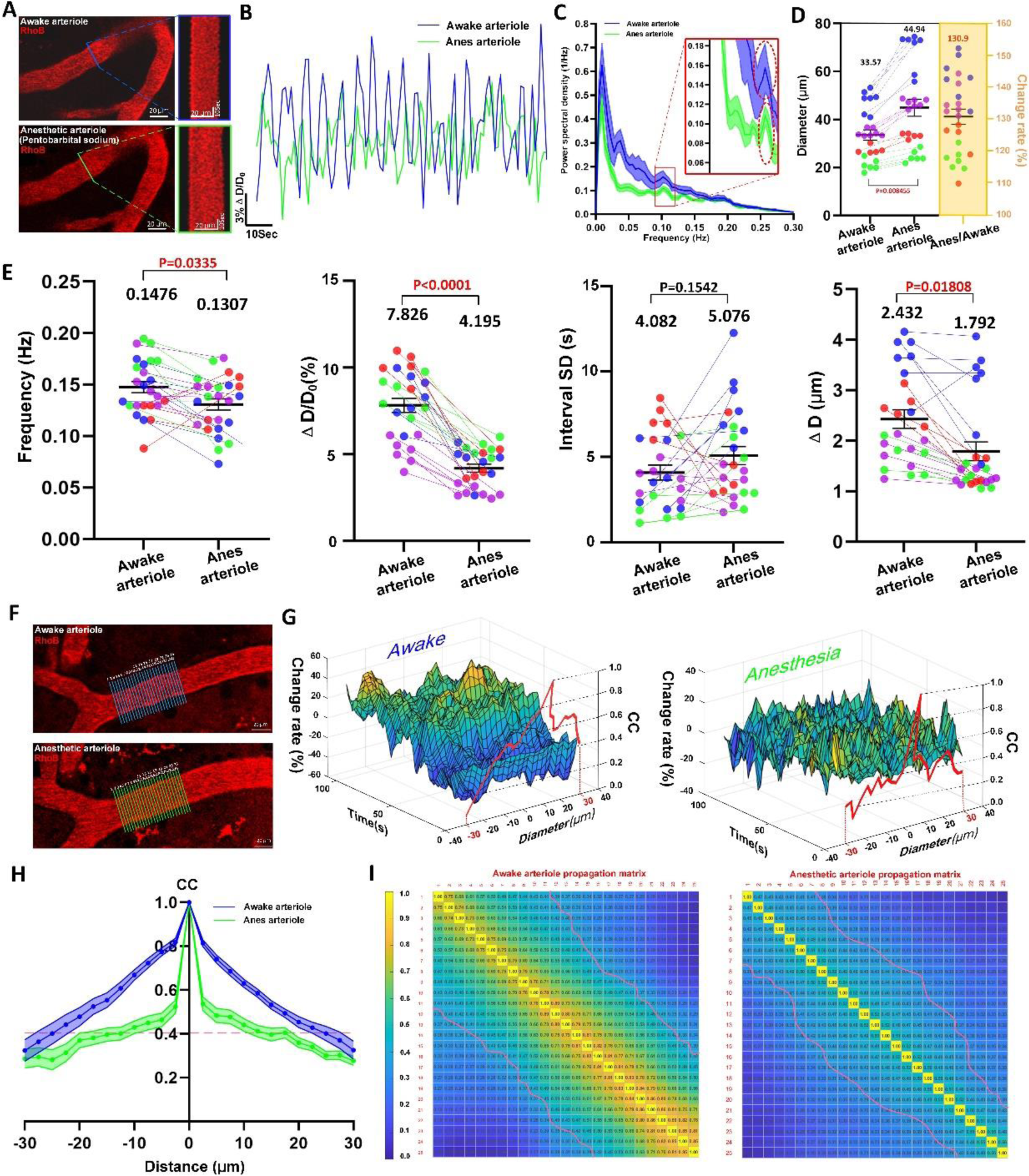
Arterioles exhibit suppressed spontaneous vasomotion and weakened vasomotion propagation index under anesthesia (A) Representative still-frame image and kymograph of same mouse cerebral arteriole in both awake and anesthetized states. (B) Representative time-lapse amplitude ratio trace of same mouse arteriole in both awake (blue line) and anesthetized (green line) states. (C) Cerebral arteriolar dilation in mice (n=25 arterioles segments in 4 mice) under anesthesia. (D) The averaged Fourier plot across same mouse arteriole diameter change in both awake (blue line) and anesthetized (green line) states (n=12 arterioles segments in 4 mice) revealed a distinct peak centered at around 0.1 Hz, while no significant peak in anesthetized mouse venule. Shaded areas represent SEM. (E) Vasomotion index comparison of same mouse arterioles before and after anesthesia (n=25 arterioles segments in 4 awake mice). The different color dots represent different mice and connecting lines represent vasomotion index change of the same arteriole in the same mouse before and after anesthesia. (F) Representative example of the diameter changes propagation dataset of same mouse cerebral arteriole in both awake and anesthetized states (G) Representative 3D mashgrid plots of arterioles before and after anesthesia. (H) The arteriolar diameter change propagation index analysis. The blue (awake) and green (anesthesia) curves show the average cross-correlation value of different distance diameter change time-profile (n=21 arteriolar diameter change propagation column in 3 mice). The magenta dash line indicates correlation coefficient is 0.4. Shaded areas represent SEM. (I) The arteriolar diameter changes propagation cross-correlation matrix (n=21 arteriolar diameter changes propagation column in 3 mice, see Methods). The magenta dash line indicates correlation coefficient is 0.4.

The Fourier spectral analysis revealed that arterioles exhibited a prominent peak at 0.1 Hz both in awake and anesthetized states (Fig. 5C). However, the arterioles performed significantly smaller amplitudes under anesthesia. Notably, the average dilation rate of the resting diameter of arterioles reached 30.90±2.384% (Fig. 5D). This may be related to the observed reduction in vasomotion amplitude revealed by the time series analysis (Fig. 5E). Moreover, the arteriole motion frequencies decreased under anesthesia (9.323% ± 4.690). The amplitude-related properties of the arterioles decreased reductions reached to 44.90±2.664% in ΔD/D0 and 25.15 ±4.739% in ΔD (Movie 6) (Fig. 5E, Table 5).

Consistent with previous reports, anesthesia significantly reduced the power of arteriolar vasomotion. Recently, arteriole wall motion was found to propagate along the vascular longitudinal axis^26^. Our study confirmed this propagation in arterioles and revealed that this propagation was robustly suppressed under the influence of anesthetics (Fig. 5F-I), suggesting a novel aspect of the side effects of anesthesia. Taken together, our findings demonstrate that anesthesia profoundly affects spontaneous vasomotion.

### Vasomotion features in different diameter ranks of vessels and relationships comparison between SMCs Ca^2+^ oscillations with vasomotions in arterioles

Next, to emphasize lumen size-based differences in our subsequent analyses, the three breakpoints were employed to categorize the vessels into four diameter ranks. These numbers corresponded to the quartiles value in normalized histograms for vessel segments^37^. For arterioles, 170 segments from sixteen mice were calculated and the ranks were defined as rank1 (38.83 < d < 56.31 µm), rank2 (32.18 < d < 38.83 µm), rank3 (23.05 < d < 32.18 µm), and rank4 (17.80 < d < 23.05 µm).

We then characterized differences in vasomotion indices arterioles of different ranks. After a comprehensive analysis, representative kymographs and ΔD/D0 time series indicate that smaller vessels in arteriolar networks exhibited more pronounced vasomotion (Fig. 6A-C, Table 6). We found no differences in the frequency or interval SD among arterioles with different ranks. Intriguingly, the ΔD values of the arterioles were consistently approximately 2 µm, regardless of ranks.

**Figure. 6.**
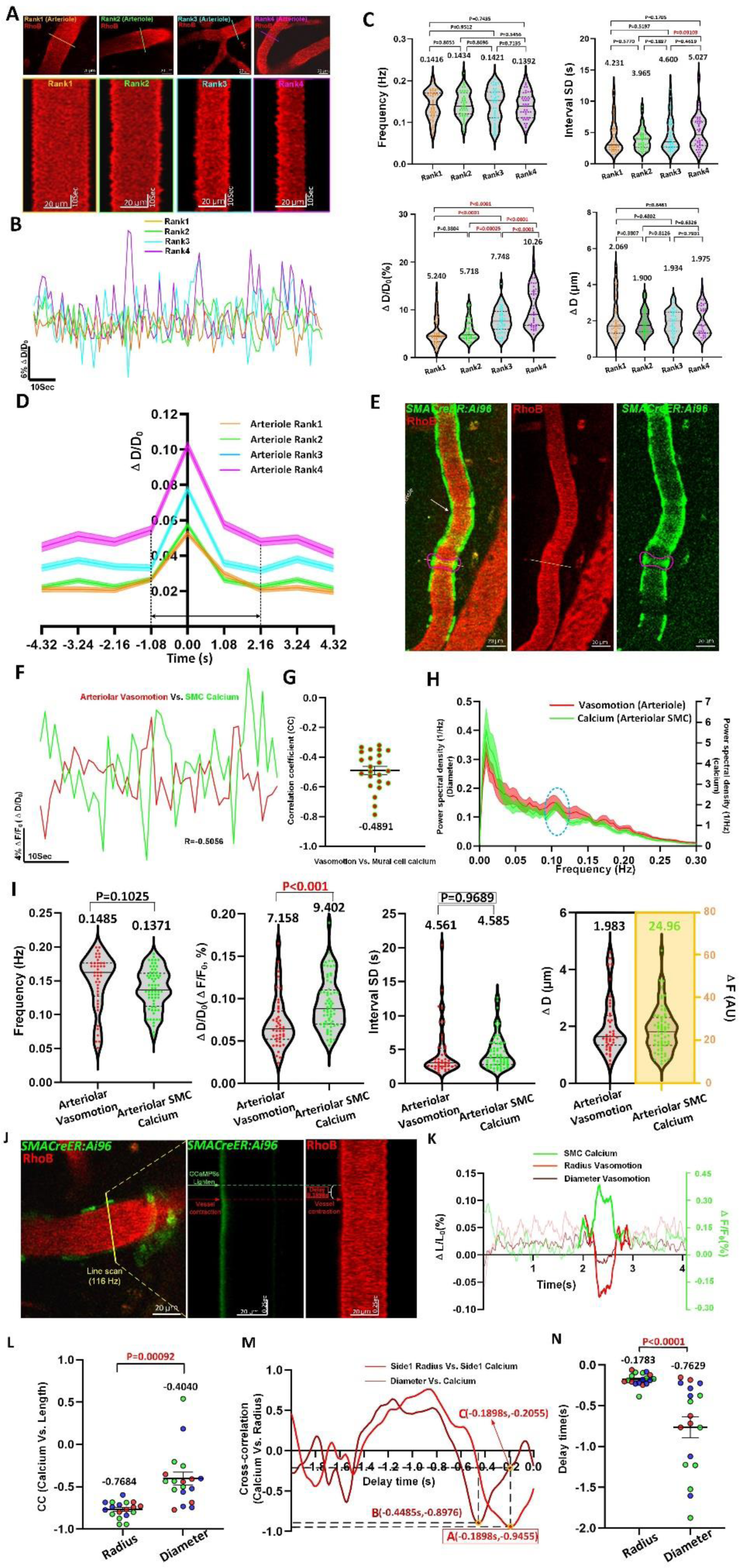
Vasomotion features in different diameter ranks of vessels and relationships comparison between SMCs Ca^2+^ oscillations with vasomotions in arterioles. (A) Representative still-frame image and kymograph of different rank cerebral arterioles. The 170 arteriole segments collected from 16 awake mice were ranked by Quartile distribution of diameter. For arterioles, Rank1: 38.85-56.31 μm (n=42 arterioles segments in 11 awake mice), Rank2: 32.18-38.85 μm (n=43 arterioles segments in 8 awake mice), Rank3: 23.05-32.18μm (n=43 arterioles segments in 10 awake mice), Rank 4: 17.80-23.05 μm (n=42 arterioles segments in 7 awake mice). (B) Comparison of representative time-series amplitude changes trace at four diameter rank levels of arterioles. (C) Vasomotion index comparison of arterioles at four diameter ranks (n=170 arterioles segments in 16 awake mice). (D) The averaged single vasomotion events plot comparison of arteriole at four diameter ranks. For arterioles, Rank1: n=481 vasomotion events in 7 awake mice, Rank2: n=486 vasomotion events in 5 awake mice, Rank3: n=510 vasomotion events in 6 awake mice, Rank4: n=493 vasomotion events in 5 awake mice. (E) Representative still-frame images including vessels vasomotion and GCaMP6s calcium fluorescence intensity changes of corresponding mural cell (purple mural cell shape curve). (F) Representative time-series mural cell calcium amplitude ratio (ΔF/F0, green solid line) and vessels vasomotion amplitude ratio (ΔD/D0, red solid line) change traces of awake mouse, and the CC comparison of two time series curves. (G) The quantification of CC values between arteriolar vasomotion and SMCs calcium (n=23 pairs in 5 awake mice). (H) The averaged Fourier plot across mouse arterioles SMCs calcium signal change and arteriolar diameter change in awake state revealed a distinct peak centered at around 0.1 Hz (n=38 SMCs in 6 awake mice Vs. n=21 arterioles segments in 6 awake mice). Shaded areas represent SEM. (I) Comparison between vasomotion index and calcium index (n=64 SMCs in 5 awake mice) (J) Representative still-frame image and kymograph of cerebral arteriole using high frequency line scan. (K) Representative time series of calcium fluorescence ratio ΔF/F0, diameter amplitude ratio ΔD/D0 and radius amplitude ratio ΔR/R0 for one arteriole (J) in the field. (L) Left shows quantification averaged minimum cross correlation values comparison across arteriolar radius with SMC calcium, Right shows quantification averaged CC values when arteriolar diameter amplitude ratio with SMC calcium amplitude ratio lives up to time delay value corresponding to arteriolar radius and SMC calcium amplitude ratio (n=18 arterioles segments in 3 awake mice, different color dots represent different mouse). (M) The cross-correlation to determine the delay time between time series of calcium amplitude ratio and radius or diameter amplitude ratio used for the example in (J). The radius lags electrical activity (minimum of cross correlation) by 0.1898 s and the diameter lags electrical activity by 0.4485 s. (N) The quantification averaged delay time comparison across arteriolar radius or diameter with SMC calcium (n=18 arterioles segments in 3 awake mice, different color dots represent different mouse).

Additionally, we isolated and aligned individual vasomotion events (Fig. 6D). To our knowledge, this is the first time that the kinetics of single vasomotion events have been revealed. Our findings indicate that each vasomotion event cycle required approximately one second to dilate and two seconds to contract. Ideally, as 0.1-Hz rhythmicity implies one event every 10 seconds, our findings suggest that the vessel wall remains still during the remaining six to seven seconds in each cycle. These analyses represent a comprehensive characterization of arteriolar networks.

To determine the role of Ca^2+^ oscillations in SMC in driving spontaneous vasomotion, we used awake *SMACreER:Ai96* mice. Intraluminal injection of rhodamine into these mice allowed simultaneous measurements of Ca^2+^ oscillations and lumen diameter dynamics (Fig. 6E-F). The same baseline smoothing method was used to determine the baseline Ca^2+^ signal. Consistent with previous studies^6, 21^, the SMC Ca^2+^ dynamics were inversely correlated with arteriolar vasomotion (-0.4891±0.02710, p<0.05) (Fig. 6G, Table 7, Movie 7). If Ca^2+^ dynamics are the driver of vasomotion, Ca^2+^ oscillations should have a 0.1 Hz peak in spectral domain.

Indeed, we identified this peak in arteriole SMCs (Fig. 6H). Furthermore, there is no differences in the frequency and interval SD between Ca^2+^ oscillations and vasomotion (Fig. 6I, Table 8).

Next, we employed a high-speed line scanning approach to determine how early SMC Ca^2+^ activity leads to vasocontraction (Fig. 6J). The time series of changes in diameter, radius, and Ca^2+^ activity revealed a clearly mirrored pattern between vasomotor activity and Ca^2+^ oscillations when a vasomotion event occurs (Fig. 6K). The CC value calculated between the changes in diameter and GCaMP6s fluorescence intensity was -0.4040±0.07691 (Fig. 6L), which is close to previous findings^6^. Notably, we found a significantly higher negative correlation of -0.7684±0.02199 when we used radius parameter to calculate the CC value. Furthermore, we found that SMC Ca^2+^ activity led to vasocontraction in 178.3±18.03 ms, which was nearly one-quarter of the time calculated using the diameter (Fig. 6M-N). Thus, we conclude that although the diameter and radius indices are interchangeable in most vasomotion analyses, when observing actual time delay result between SMC Ca^2+^ oscillations and vasomotion (Fig. 6J), radius-based vasomotion analysis is more precise.

### During ischemic stroke, arteriole vasomotion impairment is an early pathology that persists for a long time

We implemented two hours filament occlusion of the MCA and performed in vivo repeated imaging of the same MCA branches before occlusion, during occlusion, and 1 or 14 days after recanalization. We observed Ca^2+^ overload in SMCs, with signal amplitudes four times higher than basal levels (Fig. 7A-B). In addition, obvious nonisotropic movements between two sides of the vascular wall were observed in the kymograph (Fig. 7A, Movie 8). These Ca^2+^ elevations were immediately followed by strong SMC constrictions. This led to a reduction in the resting diameter from 32.82 µm to 12.29 µm within 22 seconds and in the sequential fast and slow recovery phases, and recovery to baseline was not achieved within 200 seconds (Fig. 7A-B). These data suggest that SMC Ca^2+^ overload is likely the central pathological driver of arteriolar constriction.

**Figure. 7.**
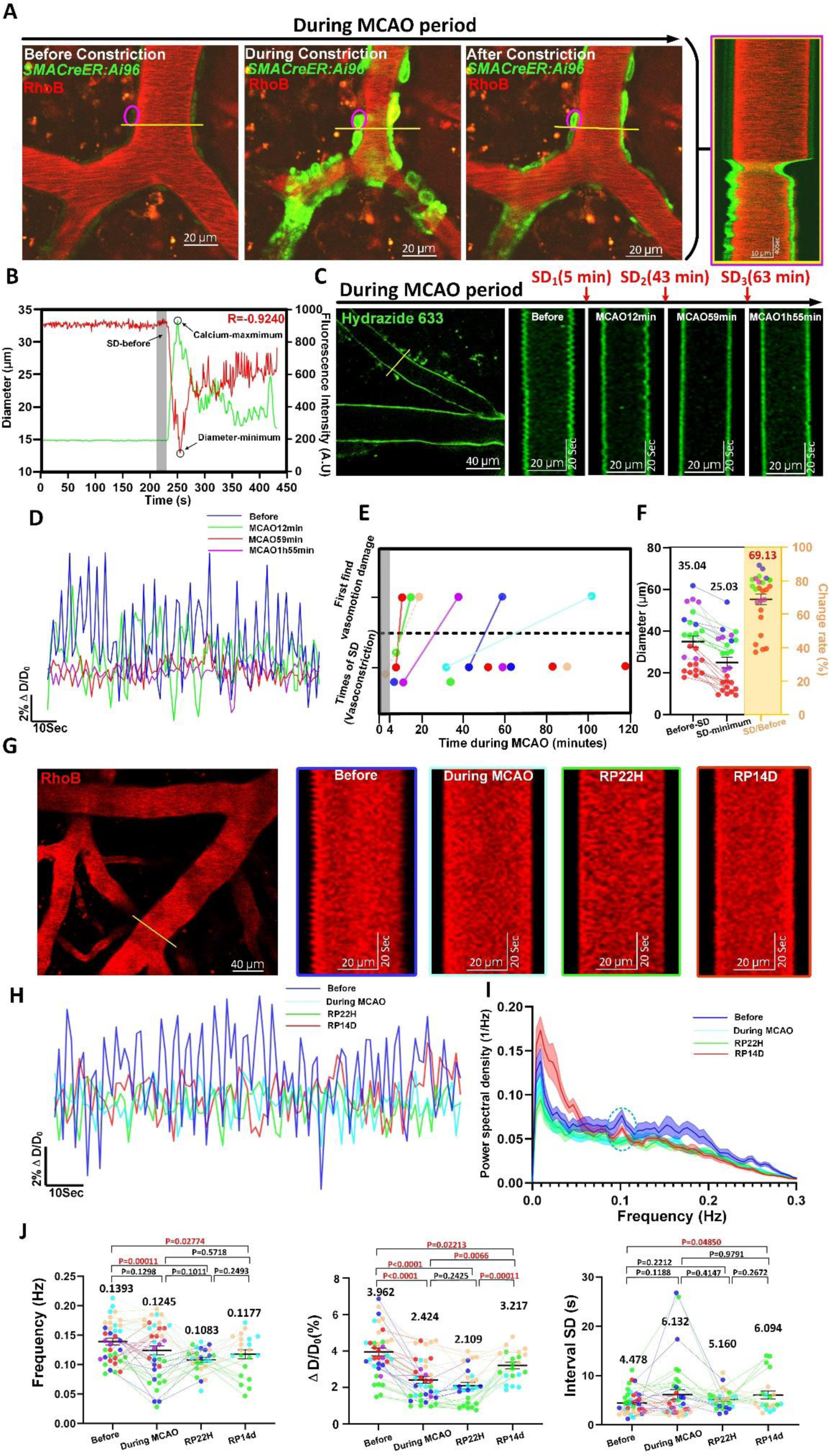
Vasomotion damage immediately takes place after the vasoconstriction during the initial minutes of occlusion and lasts long despite successful recanalization. (A) Time-series sequence and kymograph showing both GCaMP6s calcium changes in a single SMC and cerebral arteriole diameter change undergoing spread depolarization (SD) vasoconstriction during MCAO period. The yellow line represents the reslice position. (B) Representative time-series calcium signal and diameter change trace of cerebral arteriole in anesthetized mouse during MCAO period. Shaded areas represent average diameter of 15 frames (16.2s) before SD. (C) Time-series sequence and kymograph showing the degree of vasomotion damage during the occurrence of SD in MCAO period. (D) Representative time-series amplitude changes trace of same cerebral arteriole during different stroke periods. (E) Top shows the first time vasomotion damage observed during MCAO period (N=6 anesthetic mice). Bottom shows times of SD within 4-120 minutes of MCAO (n=14 in 6 anesthetic mice). The gray column represents the time (4 minutes) taken for the mouse to 2P platform for imaging after MCAO surgery. The connecting solid lines represent time interval between vasomotion damage happening and first SD occurred during in vivo imaging recorded (the dashed line is estimated time interval). (F) Cerebral arteriolar vasoconstriction in mice (n=25 arterioles segments in 6 mice) under anesthesia. (G) Representative still-frame image and kymographs of same cerebral arteriole during different stroke periods. (H) Comparison of representative time-series amplitude changes trace of same arteriole at four periods (Before, During MCAO, RP22H and RP14D). (I) The averaged Fourier plot across same mouse cerebral arteriole diameter change at four stages (n=26 arterioles segments in 6 mice of before stroke stage, n=26 arterioles segments in 6 mice of during MCAO stage, n=20 arterioles segments in 4 mice of RP22H stage and n=17 arterioles segments in 3 mice of RP14D stage) revealed a distinct peak centered at around 0.1 Hz of stage before and RP14D, while no significant peaks of stage during MCAO and RP22H. Shaded areas represent SEM. (J) Vasomotion index comparison of same arterioles at four stages (n=37 arterioles segments in 6 anesthetic mice of before stroke stage, n=37 arterioles segments in 6 mice of MCAO damage stage, n=27 arterioles segments in 4 mice of RP22H stage and n=21 arterioles segments in 3 mice). Due to the partial death of stroke mice and decrease in imaging effect of the cranial window, the number of kinetics characteristics detecting mice decreases over time. The different color dots represent different mice and connecting lines represent diameter or vasomotion index change of the same arteriole in the different stages during stroke.

Multiple episodes of SD-induced vasoconstriction occur during ischemia^6, 42^, and we hypothesized that these early constrictions gradually lead to damaged vasomotion. For instance, we observed only slight and moderate deterioration 7 min after the first episode (MCAO 12 min) and 16 min after the second episode (MCAO 59 min), respectively, while we detected nearly no vasomotor activity 52 min after the third episode (MCAO 1h55 min) (Fig. 7C-D). In total, we examined six mice, each of which displayed 1–4 episodes and an average reduction in the resting diameter of 30.87 ± 3.200% (Fig. 7E-F). Notably, we consistently observed impaired vasomotion following the initial vasoconstriction in five out of six mice (Fig. 7E). We may have missed the first vasoconstriction in the mouse represented by the peach-colored dots due to the blank four-minute period required to settle the animal under the microscope. These findings indicate that spontaneous vasomotion impairment follows pathological arteriolar constriction driven by SMC Ca^2+^ overload during early ischemia. and repeated SMC Ca^2+^ overload has an immediate and accumulative detrimental impact on basal vasomotion.

Next, to determine the extent of vasomotion damage, we examined vasomotor activity at the same MCA branches position during different stroke periods (Fig. 7G). It revealed that arterioles transitioned to an inert state (during MCAO) and showed very mild recovery (recanalization 14 days) (Fig. 7H-I). In addition, the vasomotion indices of frequency and ΔD/D0 reached their lowest values approximately one day after recanalization and recovered only slightly 14 days after recanalization (Fig. 7J, Table 9). Our findings suggest that long-term vasomotion damage is a previously unrecognized vascular pathology associated with stroke that may impact postischemic blood flow. Consequently, targeting the early dysfunction of myogenic spontaneous vasomotion could be a novel therapeutic approach for preventing reperfusion deficits.

## Discussion

In this study, we first examined spontaneous vasomotion in arteriolar networks in the mouse brain under various conditions, such as ex vivo, head-fixed awake, anesthesia, and focal ischemia. By adopting a time-rolling baseline method to investigate vasomotion (Fig. 1), we overcame long-standing challenges: we isolated single vasomotion events and identified arteriolar vasomotion of different ranks. Then, we comprehensively and quantitatively characterized thirteen aspects of vasomotion, including kymographs, the time series amplitude ratio, spectrum density, vasomotion indices (frequency, interval SD, ΔD, ΔD/D0 and resting average diameter), single vasomotion event kinetics, CC and DTW relationships, vasomotion propagation, and the time latency between SMC Ca^2+^ activity and vasomotion. For the first time, we considered the radius parameter in vasomotion, identifying asynchronous movement of the vascular wall at transverse cross-sections (Fig. 4). Importantly, we identified a novel vascular pathology in which arterioles transitioned from a vasoactive to a vaso-inactive status following repeated SD-induced vasoconstrictions during the super acute ischemia period, with more SD episodes leading to more severe vasomotion damage (Fig. 7). Vasomotion can serve as a valuable indicator to assess the extent of vascular damage under ischemic stroke. These quantitative descriptions of spontaneous vasomotor activity improve our understanding of functional features of cerebral vasculature.

Our findings in ex vivo brains, in which presumably no neural activity was present after being in ACSF for two hours, support the role of the intrinsic contractility of SMCs in generating arteriolar vasomotion through self-generated movement^22^ (Fig. 2). Then the 3.24-second vasomotion events measured for arterioles are consistent with recent findings on the average duration of the coinciding SMC Ca^2+^ peak^21^. This consistency suggests that single events were reliably isolated in the present study and that SMCs Ca^2+^ fluctuations influence vasomotion kinetics.

Furthermore, exploring the relationship between cerebrospinal fluid (CSF) flow and vasomotion is of utmost importance. From a dorsal view, CSF flow speed differs along the two sides of pial arteriole wall^43^. Together with previous work suggesting that vasomotion promotes CSF flow, our findings on asymmetric movements of arteriolar walls (Fig. 4) may explain the asymmetry observed in CSF flow. Additionally, different vasomotion amplitudes across different ranks in arteriolar network (Fig. 6) suggest potentially different potencies in CSF flow, which might lead to different spatial clearing rates of β-amyloid (Aβ). Indeed, studies on both AD patients and AD model animals have reported that Aβ deposits do exhibit spatial differences related to the size of arteries^44–46^. These results suggest that strategies to specifically manipulate and target arteriole segments should be developed in future work.

A limitation of this study is the low scanning rate of 0.926 Hz when using a 512 × 512 FOV; as a result, we could obtain only the low-frequency domain in the vasomotor waves, which may lead to missing information. In the future, when using conventional TPLSM to study vasomotion, small FOVs, such as 128 × 128, should be selected to capture higher frequency domains.

This study confirmed our recent findings that pial arterioles transition to an inert state during the first 24 hours after recanalization^27^. However, how early vasomotion damage is induced and how long vasomotion damage lasts have not yet been established. We addressed these important questions by investigating the super acute period and a longer reperfusion period (RP 14 days) (Fig. 7) and found that the number of SDs appeared to have a cumulative effect on the extent of vasomotion damage (Fig. 7C). Therefore, we suspect that increased transient Ca^2+^ activity can cause more severe damage to molecular machines that are responsible for cyclic SMC contractility. Prevention of SMC Ca^2+^ overload could represent a novel therapeutic goal to prevent delayed damage to vascular function.

In conclusion, comprehensively quantifying vasomotion is essential for understanding cerebral vascular networks, assessing vascular healthy and exploring vasomotion therapeutic interventions. This quantitative approach enhances the knowledge of vessel dynamics, and ultimately contributes to the development of improved treatments for various brain disorders such as stroke, Alzheimer’s disease and vascular dementia.

## Supporting information

FigureS1-S5, tables, movie legends and algorithm formulas

## Acknowledgements

We are very grateful to Drs. Xu-zhao Li, Drs. Dong-dong Zhang and Mr. Jia-yu Ruan for constructive suggestions during this study.

## Author contribution statements

Yi-yi Zhang, Conceptualization, Resources, Data curation, Software, Formal analysis, Validation, Investigation, Visualization, Methodology, Writing – original draft; Jin-ze Li, Resources, Conceptualization, Data curation, Formal analysis, Validation, Investigation, Writing—review and editing; Hui-qi Xie, Resources, Conceptualization, Formal analysis; Yu- xiao Jin, Validation, Investigation; Wen-tao Wang, Investigation, Methodology; Bing-rui Zhao, Validation, Investigation; Jie-min Jia, Conceptualization, Funding acquisition, Project administration, Resources, Supervision, Writing review and editing.

