## Supplementary material for "High-resolution vasomotion analysis reveals novel arteriole physiological features and progressive modulation of cerebral vascular networks by stroke": FigureS1-S5, tables, movie legends and algorithm formulas

#### Supplementary figures

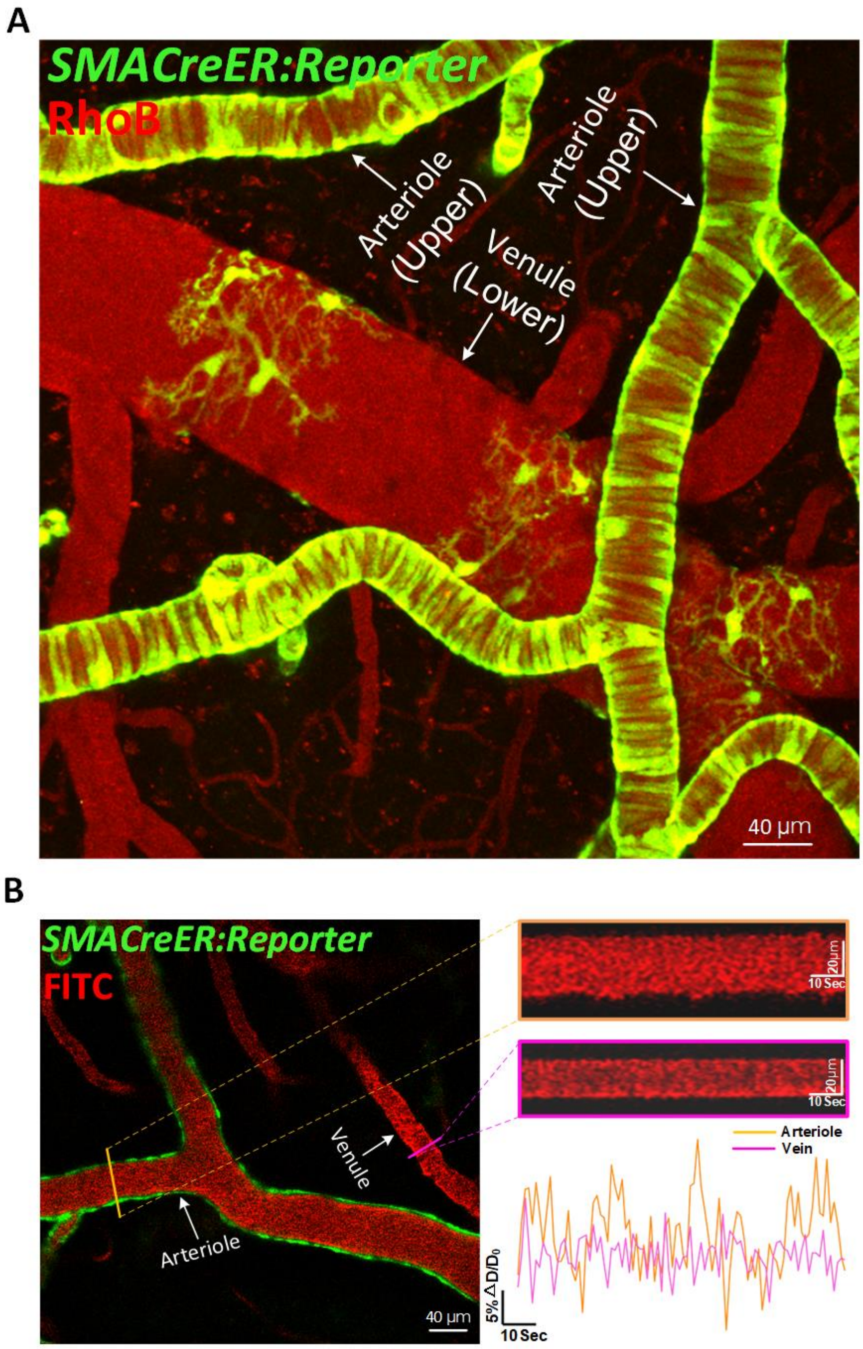


**Figure.S1 Classification of arterioles and venules and myogenic spontaneous vasomotion difference between cerebral arteriole and venule in awake mouse. (A) Maximal intensity projection (MIP) of the cerebral pial vessels, including the arterioles and venules in *SMACreER: Ai47* mice injected with RhoB intravenously under 2P. The arterioles and venules can be distinguished by the shape of the mural cell warpped**

**around them. (B) Representative still-frame image and kymograph of cerebral arteriole and venule. The yellow solid line represents the reslice position of arteriole. The purple solid line represents the reslice position of venule. Right bottom shows representative time series amplitude changes ratio of cerebral arteriole (yellow solid line) and venule (purple solid line).**

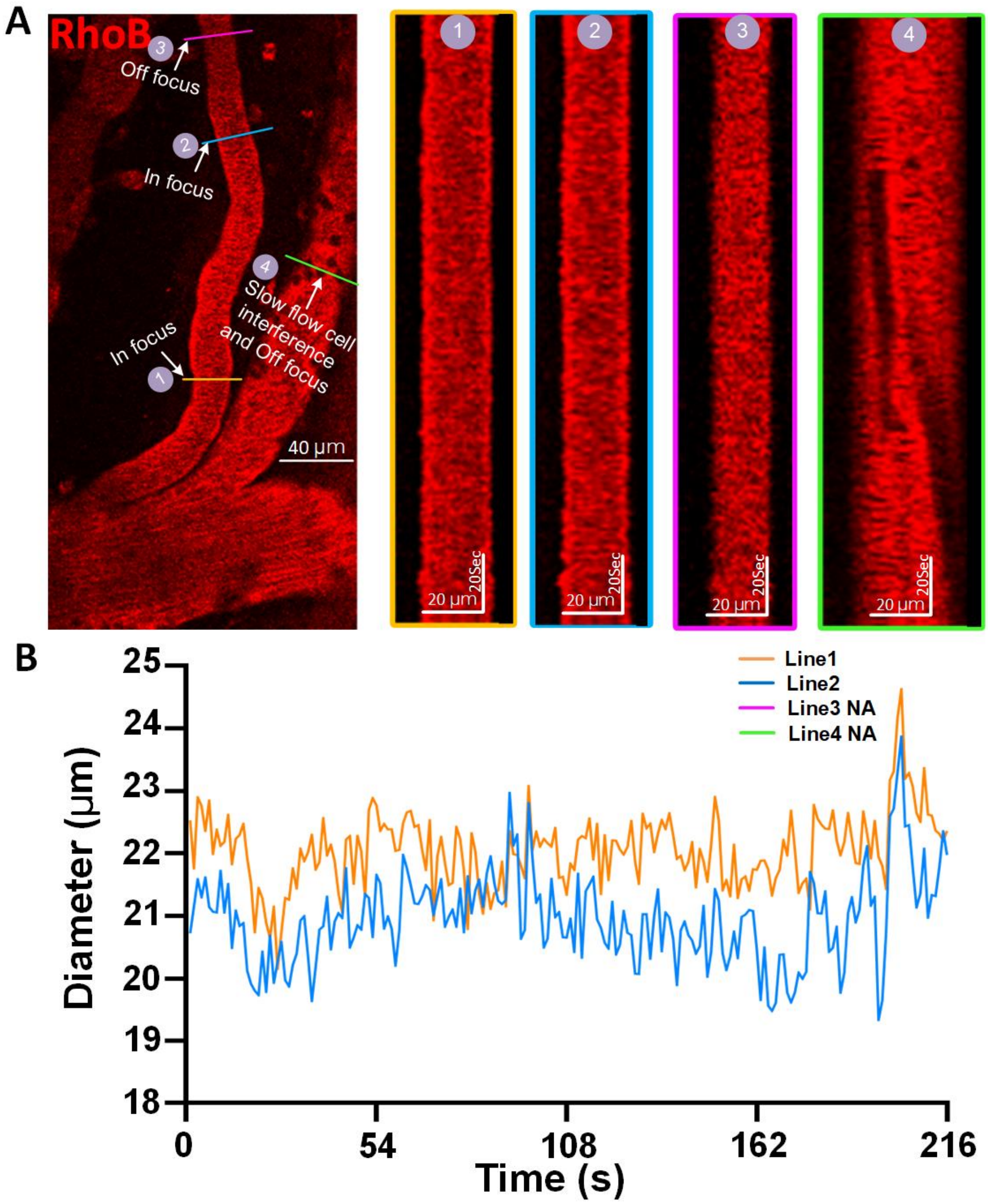


**Figure.S2 Selection of vasomotion analysis data. (A) Representative still-frame image and kymograph of different arteriole segments. The yellow, blue solid line represents the line reslice position of in-focus data and the purple and green line represents the line reslice position of out-focus data. (B) Representative time-series diameter change trace of mouse arteriole (line 1-2) in awake states. NA represents the out-focus data (line 3-4) could not be quantified as diameter change trace due to unclear boundaries.**

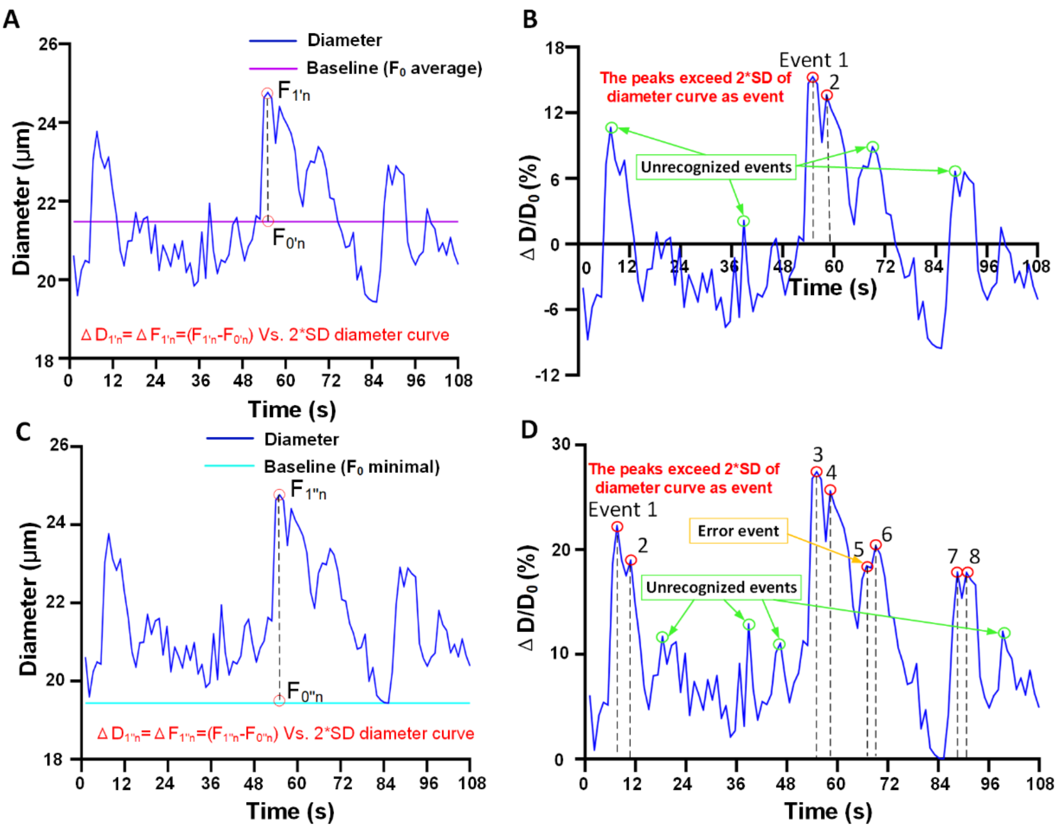


**Figure.S3 Extracting the baseline through $F_0$ average and $F_0$ minimal in inaccurate events identification. (A) Representative time-series diameter changes trace and baseline of $F_0$ average (purple line, the mean of diameter change trace). The event was judged by whether the $F_{1'n}$- $F_{0'n}$ was exceed the double SD of diameter change trace. (B) Representative time-series amplitude ratio ($\Delta D/D_0$) changes trace over baseline $F_0$ average and events annotation. The green dots represent some unrecognized but obvious events and the red dots represent the partially marked events. (C) Representative time-series diameter changes trace and baseline of $F_0$ minimal (cyan line, the minimum value of diameter change trace). (D) Representative time-series amplitude ratio ($\Delta D/D_0$) changes trace over baseline $F_0$ minimal and events annotation. The green dots represent some unrecognized but obvious events and the red dots represent the partially marked events but contain misidentification event.**

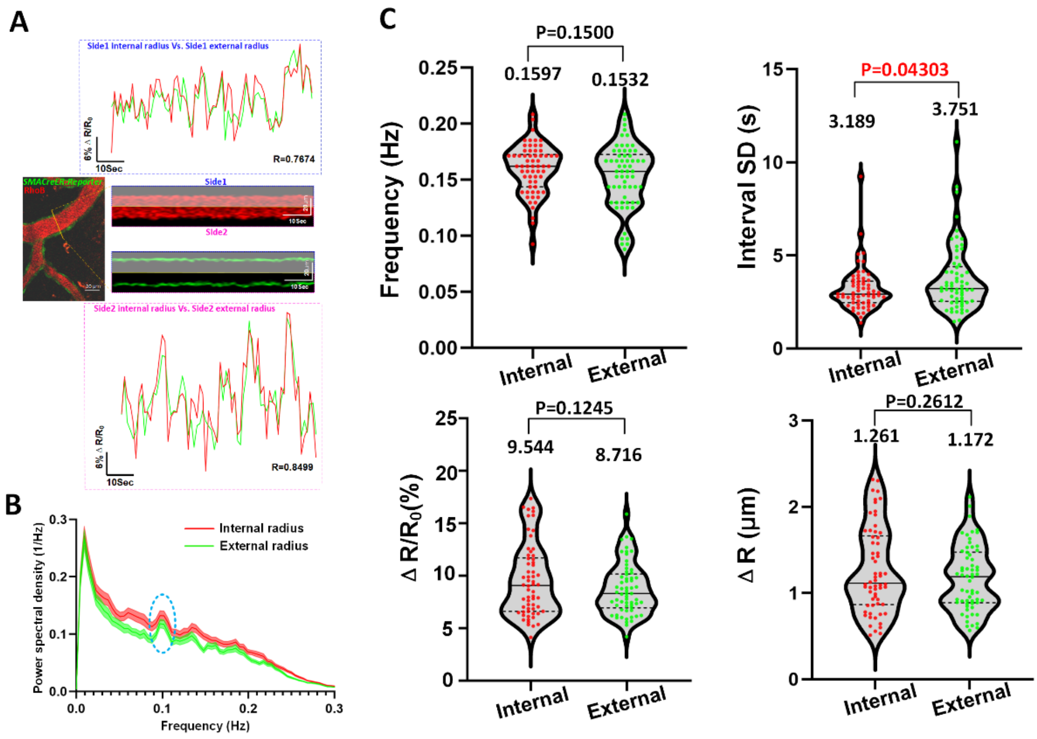


**Figure.S4 All dynamic changes in internal and external vascular radius can be used to characterize spontaneous vasomotion. (A) Representative still-frame images and kymographs of the same cerebral arteriole internal (red) and external (green) radius and corresponding CC value of time-series external-internal diameter amplitude ratio $\Delta R/R_0$ change curves. The yellow solid line represents the reslice position. (B) The averaged Fourier plot across awake mouse cerebral internal and external radius change of same arterioles (n=62 arterioles segments in 8 awake mice) revealed a broad range of ultra-low frequencies (<0.3Hz), with a distinct peak centered at around 0.1 Hz. Shaded areas represent SEM. (C) Vasomotion index analysis of cerebral arteriole between internal and external radius (n=62 arterioles segments in 8 awake mice). Top left represents the index of frequency, top right represents the SD of the adjacent vasomotion event interval, bottom left represents the index of amplitude ratio $\Delta R/R_0$ and bottom right represents index of absolute amplitude $\Delta R$. The reporter mice used in our study included Ai14, Ai47,**

**and Ai96 mice which external diameter are uniformly labeled as green color and internal diameter are labeled as red color.**

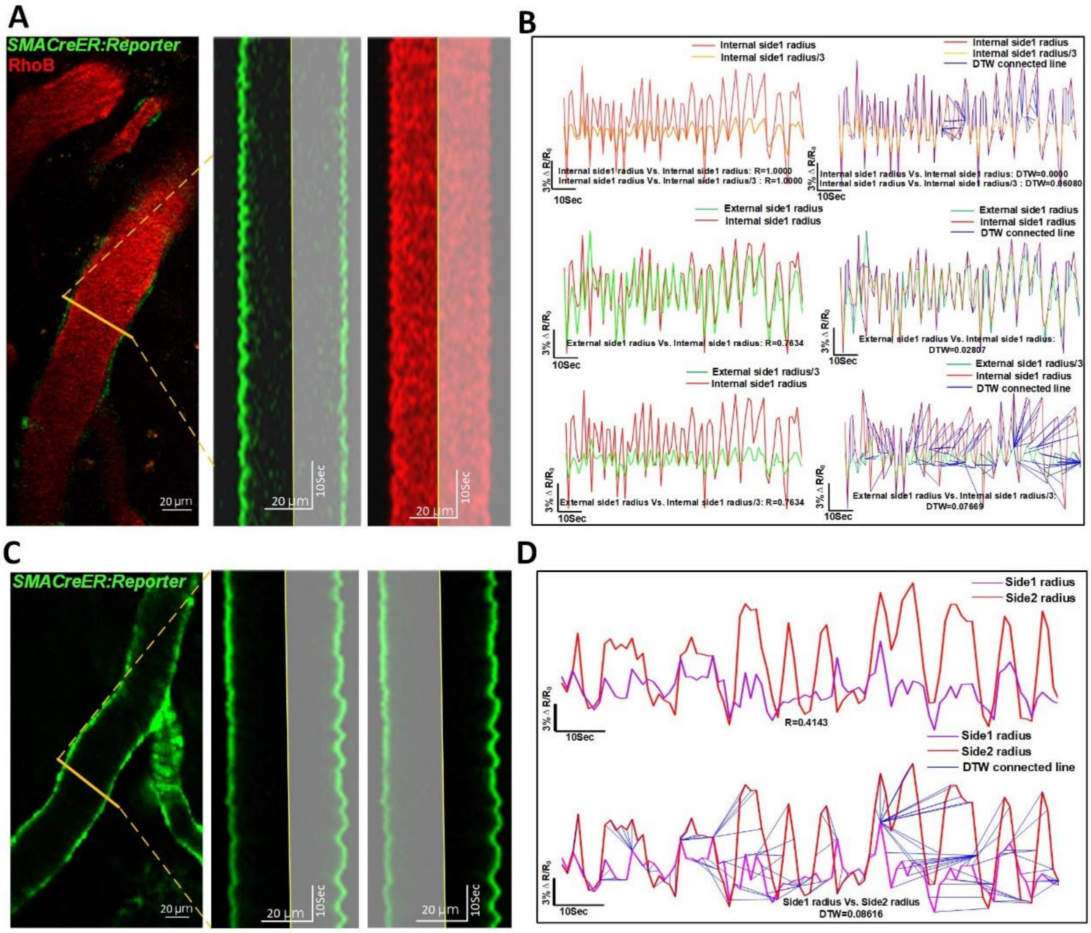


**Figure.S5 Application of Dynamic Time Warping (DTW) algorithm in comparing curve difference between internal radius and external radius amplitude ratio $\Delta R/R_0$, as well as the curve difference between side1 and side2 radius amplitude ratio. (A) Left shows representative in vivo 2P microscopy images of cerebral arteriole through a cranial window in an awake head-fixed mouse. Middle shows the kymographs of external side1 radius and right shows the kymographs of internal side1 radius. Shaded areas indicate nonrepresentative radius of the other side. (B) Top left shows time series of internal side1 radius $(\Delta R/R_0)_i$ (red solid line) and internal side1 radius $(\Delta R/R_0)_i/3$ (yellow solid line) amplitude ratio change curves and the corresponding CC value comparison. Middle left shows time series of external side1 radius $(\Delta R/R_0)_e$ (green solid line) and internal side1**

**radius $(\Delta R/R_0)_i$ (red solid line) amplitude ratio change curves and the corresponding CC value comparison. Bottom left shows time series of external left radius $(\Delta R/R_0)_e/3$ and internal side1 radius $(\Delta R/R_0)_i$ amplitude ratio change curves and the corresponding CC value comparison. The CC value remains unchanged if only the proportional scaling of amplitude is employed. Top right shows the DTW value of figure top right, Middle right shows the DTW value of figure middle left and Bottom right shows the DTW value of figure bottom left. There are high DTW value (large curves difference) of Bottom right compare to Top right and middle right. The CC value did not reflect similarity in the amplitudes of the curves, as demonstrated by the constant R value as the amplitude decreased, while the DTW score increased with decreasing amplitude. Similarly, when comparing the mathematically scaled-down external radius with the internal radius, the R value remained constant while the DTW score increased. (C) Representative still-frame image and kymograph of cerebral arteriole external side1 and radius external side2 radius. (D) Top shows time series of external side1 radius and external side2 radius amplitude ratio change curves and the corresponding CC value comparison. Bottom shows the DTW value of top.**

### Supplementary tables

| Awake mice | Frequency (Hz) | | Interval SD (s) | | Amplitude ratio ($\Delta D/D_0$,%) | | Absolute amplitude ($\Delta D$, μm) | | Number | |
|---|---|---|---|---|---|---|---|---|---|---|
| | Awake | Ex vivo | Awake | Ex vivo | Awake | Ex vivo | Awake | Ex vivo | Awake | Ex vivo |
| Arteriole (diameter) | 0.1471± 0.005107 | 0.07851± 0.004232 | 4.094± 0.4286 | 7.125± 0.7546 | 7.202± 0.4442 | 1.972± 0.1610 | 1.962± 0.1286 | 1.096± 0.07489 | N=4, n=38 | N=4, n=30 |

**Table. 1 Vasomotion index between awake mouse arterioles and ex vivo brain arterioles**

| Awake mice | Frequency (Hz) | | Interval SD (s) | | Amplitude ratio ($\Delta D/D_0$,%) | | Absolute amplitude ($\Delta D$, μm) | | Correlation coefficient | Number | |
|---|---|---|---|---|---|---|---|---|---|---|---|
| | Internal | External | Internal | External | Internal | External | Internal | External | Internal Vs. External | Internal | External |
| Arteriole (diameter) | 0.1543± 0.004360 | 0.1455± 0.006184 | 3.282± 0.1887 | 4.133± 0.4155 | 7.449± 0.5309 | 6.401± 0.3703 | 1.876± 0.1219 | 1.613± 0.09397 | 0.6855±0.01542 | N=8, n=31 | N=8, n=31 |
| Arteriole (radius) | 0.1597± 0.002786 | 0.1532± 0.003579 | 3.189± 0.1496 | 3.751± 0.2309 | 9.544± 0.4363 | 8.716± 0.3101 | 1.261± 0.06318 | 1.172± 0.04797 | 0.7074±0.01140 | N=8, n=62 | N=8, n=62 |

**Table. 2 Vasomotion index and correlation coefficient in arteriolar internal and external diameter, internal and external radius.**

| Awake mice | Correlation coefficient | | | Number |
|---|---|---|---|---|
| Arteriole | External radius Vs. Internal radius | | | N=8, n=62 |
| | 0.7074±0.01140 | | | |
| Arteriole | Diameter Vs. Side1 radius | Diameter Vs. Side2 radius | Side1 radius Vs. Side2 radius | N=9, n=62 |
| | 0.7223±0.01981 | 0.7455±0.01820 | 0.1729±0.04281 | |

**Table. 3 Correlation coefficient between awake mouse arteriolar diameter and radius.**

| Awake mice | Frequency (Hz) | Interval SD (s) | Amplitude ratio ($\Delta D/D_0$,%) | Absolute amplitude ($\Delta D$, μm) | Number |
|---|---|---|---|---|---|
| Arteriole (diameter) | 0.1392±0.003213 | 4.571±0.2506 | 6.785±0.3718 | 0.8634±1.867 | N=9, n=89 |
| Arteriole (side1 radius) | 0.1435±0.003482 | 4.593±0.3502 | 8.695±0.5213 | 1.175±0.0424 | N=9, n=89 |
| Arteriole (side2 radius) | 0.1423±0.003208 | 4.308±0.2883 | 9.115±0.5013 | 1.253±0.04463 | N=9, n=89 |

**Table. 4 Vasomotion index between awake mouse arteriolar diameter and radius.**

| | Frequency (Hz) | | Interval SD (s) | | Amplitude ratio ($\Delta D/D_0$,%) | | Absolute amplitude ($\Delta D$, μm) | | Average diameter (μm) | | Number | |
|---|---|---|---|---|---|---|---|---|---|---|---|---|
| | Awake | Anes | Awake | Anes | Awake | Anes | Awake | Anes | Awake | Anes | Awake | Anes |
| Arteriole (diameter) | 0.1476±0.005384 | 0.1307±0.005475 | 4.082±0.4340 | 5.076±0.5323 | 7.826±0.4070 | 4.195±0.2302 | 2.432±0.1839 | 1.792±0.1862 | 33.57±2.142 | 44.94±3.543 | N=4, n=25 | |

**Table. 5 Vasomotion index between awake mouse arteriolar diameter.**

| Awake mice (diameter) | Frequency (Hz) | Interval SD (s) | Amplitude ratio ($\Delta D/D_0$,%) | Absolute amplitude ($\Delta D$, μm) | Number |
|---|---|---|---|---|---|
| Arteriole (Rank1) (38.83-56.31) | 0.1416±0.005363 | 4.231±0.4015 | 5.240±0.4189 | 2.069±0.1650 | N=10, n=42 |
| Arteriole (Rank2) (32.18-38.83) | 0.1434±0.004762 | 3.965±0.2580 | 5.718±0.3448 | 1.900±0.1000 | N=8, n=43 |
| Arteriole (Rank3) (23.05-32.18) | 0.1421±0.006075 | 4.600±0.4047 | 7.748±0.4032 | 1.934±0.09888 | N=10, n=43 |
| Arteriole (Rank4) (17.80-23.05) | 0.1392±0.004946 | 5.027±0.4123 | 10.26±0.6209 | 1.975±0.1205 | N=7, n=42 |

**Table. 6 Vasomotion index of different ranks arterioles.**

| | Correlation coefficient | Number |
|---|---|---|
| Arteriole (diameter) (Vasomotion Vs. SMC calcium) | -0.4891±0.02710 | N=5, n=23 |

**Table. 7 Correlation coefficient between awake mouse arterioles and SMCs calcium.**

| Awake mice (diameter) | Frequency (Hz) | Interval SD (s) | Amplitude ratio ($\Delta D/D_0$, $\Delta F/F$, %) | Absolute amplitude ($\Delta D$ (μm), $\Delta F$ (AU)) | Number |
|---|---|---|---|---|---|
| Vasomotion | 0.1485±0.006286 | 4.561±0.5837 | 7.202±0.4442 | 1.983±0.1543 | N=5, n=42 |
| Calcium | 0.1371±0.003788 | 4.585±0.3088 | 9.402±0.3830 | 24.96±1.311 | N=5, n=64 |

**Table. 8 Vasomotion index between awake mouse arterioles and SMCs calcium.**

| Anesthetic mice | Frequency (Hz) | Interval SD (s) | Amplitude ratio ($\Delta D/D_0$, %) | Number |
|---|---|---|---|---|
| Before | 0.1393±0.005710 | 4.478±0.3877 | 3.962±0.2145 | N=6, n=37 |
| During MCAO | 0.1245±0.007782 | 6.132±0.9731 | 2.424±0.1848 | N=6, n=37 |
| RP22h | 0.1083±0.003973 | 5.160±0.3669 | 2.109±0.1829 | N=4, n=27 |
| RP14d | 0.1177±0.007663 | 6.094±0.8185 | 3.217±0.1812 | N=3, n=21 |

**Table. 9 Vasomotion index of anesthetic mouse arterioles during different stroke periods.**

**Supplementary movie legends**

**Supplementary movie 1: this movie shows an example how to classify of arterioles and venules and myogenic spontaneous vasomotion difference between cerebral arteriole and venule in awake mouse.**

**Supplementary movie 2: this movie shows an example of vasomotion dynamic in awake mouse brain**

**Supplementary movie 3: this movie shows an example of vasomotion dynamic in ex vivo mouse brain**

**Supplementary movie 4: this movie shows an example of the same cerebral arteriole internal (red) and external (green) diameter and corresponding high correlation coefficient (CC) of time-series external-internal diameter amplitude ratio ∆D/D0 changes curves.**

**Supplementary movie 5: this movie shows an example of the same cerebral arteriole internal side1 radius (purple) and internal side2 radius (red) amplitude ratio with internal and external radius amplitude ratio (internal side 2 radius Vs. external side 2 radius) as reference.**

**Supplementary movie 6: this movie shows an example of vasomotion dynamic**

**comparison in same mouse MCA branch position under awake and anesthetized states.**

**Supplementary movie 7: this movie shows an example of CC comparison between vasomotion dynamic and concomitant SMCs calcium oscillation in awake mouse.**

**Supplementary movie 8: this movie shows a pathological process of a single SMC GCaMP6s calcium oscillation and cerebral arteriole diameter change undergoing spread depolarization (SD) vasoconstriction during MCAO period.**

#### Supplementary introduction to employed algorithm formulas

##### Vessel diameter and corresponding SMC $Ca^{2+}$ signal measurement

All time-lapse pictures were analyzed in Fiji (version 2.3.0) and MATLAB (version R2021a; MathWorks) using custom-written scripts. SMC calcium, vascular diameter and radius were determined as described above from frame scan images collected at frequencies of 0.625-0.926 Hz or line-scan images collected at frequencies of 100-200 Hz. Measurements in which the awake mouse had substantially moved were excluded from further analysis. In the vessel-containing images, ROIs were drawn on several vessel segments, where pixel brightness along a line orthogonal to the blood vessel was extracted. Image stacks in Tiff format were loaded into Matlab and then underwent gaussian filtering to reduce spatial noise. The diameter of the vessel was determined as the full width at half maximum of the reslice profile[1], and the vessel mask of every time stack was created by brightness variation along horizontal pixels. The peak and valley intensity thresholds were determined visually for each image to ensure optical detection of the surface vessels. The parameters of the full width at half maximum (FWHM) were calculated by utilizing Monotone piecewise cubic interpolation (Pchip) developed by Hermite interpolation[2, 3]. The Pchip derived a necessary and sufficient condition for a cubic function to be monotonic on an interval, and these conditions are employed to develop an algorithm constructs an accurate, robust and visually pleasing piecewise cubic interpolant to data. The diameter change was obtained by sorting the diameter obtained by each time stack. In the calcium-signal-containing (*SMACreER:Ai96* and *PDGFRβCreER:Ai96*) images, regions of interest (ROIs) were drawn on several SMCs according to their outline, and the change in calcium signal under time series was reflected by changing of the average green fluorescence brightness of each SMC. The codes can be found in “1_Diameter_radius_detection” at https://github.com/JialabEleven/Vasomotion_analysis.

##### Vessel radius definition

For radius calculation, the midpoint line was generated by dividing individual diameter through two along the time series, based on the calculated FWHM. In order to reduce the influence of errors in the data and extract a robust and appropriate radius boundary, an overdetermined linear system solved by the least-square method (LSM) is employed[4]. The LSM is probably the most popular technique in statistics due to several factors. Firstly, most common estimators can be casted within this framework. For example, the mean of a distribution is the value that minimizes the sum of squared deviations of the scores. Secondly, using squares makes LSM mathematically very tractable because the Pythagorean theorem indicates that, when the error is independent of an estimated quantity, one can add the squared error and the squared estimated quantity[5]. What’s more, the mathematical tools and algorithms involved in LSM have been well studied for a relatively long time. With one variable and a linear function, the prediction of centerline is given by the following equation:

$$\hat{Q} = m + nP \quad (1)$$

where a set of $T$ pairs of observations $\{Q_i, P_i\}$ is used to find a function relating the value of the centerline $Q$ (dependent variable) to the values of a midpoint line $P$ (independent variable). $m$ and $n$ represents the intercept and the slope of the regression line, respectively. The least square method defines the estimate of these parameters as the values which minimize the sum of the squares between the measurements and the predicted values. The error $\varepsilon$ is expressed as follows:

$$\varepsilon = \sum_{i=1}^{t}(Q_i - \hat{Q}_i)^2 = \sum_{i=1}^{t}[Q_i - (m + nP_i)]^2. \quad (2)$$

The estimation uses the property that a quadratic expression reaches its minimum value when its derivatives vanish. Taking the derivative of $\varepsilon$ with respect to $m$ and $n$, setting them to zero:

$$\frac{\partial \varepsilon}{\partial m} = 2n \sum_{i=1}^{t} P_i + 2 \sum_{i}^{t} Q_i + 2tm \quad (3)$$

and

$$\frac{\partial \varepsilon}{\partial n} = 2n \sum_{i=1}^{t} {P_i}^2 + 2m \sum_{i}^{t} P_i - 2 \sum_{i}^{t} P_i Q_i. \quad (4)$$

Solving the equations (3) and 4, the following least square estimates of $m$ and $n$ as:

$$m = \bar{Q} - n\bar{P} \quad (5)$$

and

$$n = \frac{\sum_{i=1}^{t}(P_i - \bar{P})((Q_i - \bar{Q})}{\sum_{i=1}^{t}(P_i - \bar{P})^2} \quad (6)$$

where $\bar{P}$ and $\bar{Q}$ represent the means of $P$ and $Q$. LSM can be extended to more than on independent variable and to non-linear functions. The centerline is fitted by midpoint line using the least-square method above. The centerline was defined as one boundary of the side1 radius and side2 radius and the unilateral FWHM in time series as the other boundary for radius calculation, and eventually the radius was subsequently calculated. The codes can be found in “1_Diameter_radius_detection” at https://github.com/JialabEleven/Vasomotion_analysis.

**Power spectrum analysis**

Depending on the data format, power spectrum analysis was performed in MATLAB (text format) and presented the power spectral density on the y axis and frequency on the x axis according to shooting parameters. In MATLAB, Fast Fourier Transform (FFT) is employed to calculate the power spectral density of vessel diameter and radius, and mural cell calcium data. The mice vasomotion and calcium signals under awake/anesthesia were sampled at 0.625 Hz or 0.926 Hz (FFT length: 200, the overlap: 50%), and the filtered power of the ultra-low frequency spectrum (0-0.3 Hz) was shown. The codes can be found in “2_Frequency_calculation” at https://github.com/JialabEleven/Vasomotion_analysis.

**Pearson correlation coefficient and propagation index analysis**

Previously, Pearson correlation coefficient (CC) and propagation analysis was implemented not only to identify the functional connectivity between ultra-slow single-vessel bold spatiotemporal dynamics and neuronal intracellular calcium signals[6], but also to search spontaneous vasomotion propagates along pial arterioles in the awake mouse brain[7]. For this study, to investigate the interactions from one seed arteriolar vasomotion to other arteriolar vasomotion and relationship between vessels vasomotion and mural cell calcium, CC analysis was employed as an indicator of functional relationship. The definition of CC is as follows[8]:

$$\rho_{E,F} = \frac{\left[\sum_{j=1}^{n'}(E_i-\overline{E})(F_i-\overline{F})\right]/(n'-1)}{\sigma_E\sigma_F} \tag{7}$$

For functional relationship research between vessels vasomotion and mural cell calcium, n’ indicates the length of signals (100-point length, 0.926 Hz). $E$ represents the time series of vasomotion signal from vessel in one place, $F$ represents the calcium time course signal of mural cell covered around vessel at this place. The $\overline{E}$ and $\overline{F}$ represent the mean values of the vasomotion signal and calcium signal, respectively. The $\sigma_E$ and $\sigma_F$ respectively represent the standard deviation of vasomotion signal and calcium signal.

As for the interactions investigation of vasomotion at different distances, n’ indicates the length of signals (200-point length, 0.926 Hz), $E$ represents the time series of vasomotion signal from one seed arteriolar, $F$ represents the vasomotion time course signal from another arteriole. The $\overline{E}$ and $\overline{F}$ represent the mean values of the two vasomotion signals. The $\sigma_E$ and $\sigma_F$ represent the standard deviation of the two vasomotion signals. Along the arteriole, the core position is selected randomly, and the diameter change is calculated in this place with time stacks as core vasomotion. By choosing intervals of 2.5 μm, the vasomotion is calculated at each position ±30 μm from the core position. Compare the CC between each position vasomotion and the core position vasomotion, the corresponding curve can be obtained. To analyze the propagation of vasomotion under awake and anesthetized conditions in mice, multiple CC curves per vessel of various mice were summarized and averaged. In addition, the CC maps is shown as CC comparison of each vasomotion statistic position. The codes can be found in part 3 “3_Propagation_SMC” at https://github.com/JialabEleven/Vasomotion_analysis. The mashgrid 3D models, also called as the vasomotion synchronization 3D model, simultaneously presents the propagation parameters of time, distance, filtered amplitude ratio, and CC curve of vasomotion. The codes can be found in “4_3Dmodel_Synchronization” at https://github.com/JialabEleven/Vasomotion_analysis.

**Time lag estimation between SMC calcium fluctuation and vasomotion**

The datasets were simultaneously collected by high-speed line scanning at a frequency of 100-200 Hz using two-photon microscopy, resulting in a total of 2,000 serial frames for each dataset ($Ca^{2+}$ signal $U_{1-2000}$ and vasomotion signal $V_{1-2000}$). It is supposed that a cross correlation between two time series of SMC calcium dynamic $U_N$ and arteriolar segment vasomotion $V_N$

of equal length $N$ ($N \ll 2000$, selected from $U_{1-2000}$ and $V_{1-2000}$) exists obtained[9]. This vasomotion time series sliding window $V_N$ was covered at least one vasomotion event. To estimate time lags, a set of cross-correlation coefficient $T$ was conducted by sliding the $Ca^{2+}$ signal $U$ over the diameter or radius vasomotion $V$, with a forward-sliding step of one frame as interval. Firstly, the correlation coefficient $\rho_{U_d,V_d}$ between SMC calcium dynamic $U_d$ and arteriolar vasomotion $V_d$ with a time series length of $N$ starting from frame d is compared, which represents the relationship between two time series starting at same moment. Then keeping the vasomotion signal $V_d$ unchanged, calculate the correlation coefficient $\rho_{U_{d-1},V_d}$ between the $V_d$ and SMC dynamic signal $U_{d-1}$ with a times series length of $N$ at (d-1) frame. Following this pattern, the cross-correlation coefficient curve is obtained as follows,

$$T = \sum_{k=1} \rho_{U_{d-k+1},V_d}. \tag{8}$$

Based on the results of SMC calcium dynamic and arteriolar vasomotion from time-lapse images, there is a negative cross-correlation between the two[10, 11]. The time lag between SMC calcium and arteriolar vasomotion was presented as the moment when cross-correlation curve arrived the minimum correlation (most negative correlation value). The codes can be found in "5_Time_lag_estimation" at https://github.com/JialabEleven/Vasomotion_analysis.

##### Dynamic time warping analysis

Dynamic time warping (DTW) is widely used as a similarity measurement to find an optimal alignment two given (time-dependent) sequences under certain restrictions[12]. It allows a non-linear mapping of one signal to another by minimizing the distance between the two. This measurement is the best solution known for time series problems in various domains, including bioinformatics[13], medicine[14], computer vision and computer animation[15], data mining and time series clustering[16-18] and signal processing[19]. Currently, DTW has earned its popularity by being extremely efficient as the time series similarity detection which minimizes the effects of shifting and distortion in time by allowing “elastic” transformation of time series to detect similar shapes with different phases.

Because of the universal adaptation and robust advantage of DTW in time series similarity comparison, this method can be combined with CC to describe the similarity of different vasomotion motion patterns. Here, based on the conclusion that both internal and external radius and diameters can be effectively utilized to investigate the properties of spontaneous vasomotion, the control group which means the same vasomotion motion patterns was defined as the paired DTW distance comparison between time series of internal and external diameter. Then the DTW distance between time series of side1 radius and side2 radius was calculated which had significant difference compared to the control group to give vital quantitative evidence that asymmetrical motion between the two sides of the arteriolar vasomotion.

Firstly, Given two time series $X = (x_1, x_2, \dots x_M)$ and $Y = (y_1, y_2, \dots y_M)$, $M \in \mathbb{N}$ represented by the sequences of vasomotion signals (200-point length). The restriction placed on the data sequences is that they should be sampled at equidistant points in time. Intuitively the DTW distance has a small value when sequences are similar and large value if they are different. The core of the DTW is the dynamic programming algorithm which is commonly called “cost” distance function. The task of optimal alignment of the sequences becoming the task of arranging all sequence points by minimizing the “cost” distance function. Algorithm starts by building the distance matrix $D \in \mathbb{R}^{N \times M}$ representing all pairwise distances between vasomotion signal time course $X$ and $Y$. This distance matrix called the set of local cost matrix for the alignment of two sequences of $X$ and $Y$:

$$D \in \mathbb{R}^{N\times M}: c_{i,j} = \|x_i - y_j\|, i \in [1:N], j \in [1:M] \quad (9)$$

the algorithm will find the alignment path which runs through the low-cost areas. This alignment path defines the correspondence of an element $x_i \in X$ to $y_j \in Y$ following the boundary condition which assigne first and last elements of $X$ and $Y$ to each other. The total cost function associated with warping path computed with sum of local cost matrix,

$$c_p(X,Y) = \sum_{l=1}^{L} c(x_{n_l}, y_{m_l}) \quad (10)$$

where $n_l$ and $m_l$ represent the path to arrive the next target. The warping path has a minimal cost associated with alignment called the optimal warping path called $R^*$. In order to find an optimal warping path, we need to test every possible warping path between $X$ and $Y$ which could be computationally challenging due to the exponential growth of the number of optimal paths as the lengths of $X$ and $Y$ grow linearly. To get over this challenge, the dynamic programming was employed,

$$DTW(X,Y) = c_{r^*}(X,Y) = \min\{c_r(X,Y), r \in R^{N\times M}\} \quad (11)$$

where DTW distance means the similarity description between two vessels time series motion curves (200-point length). The codes can be found in “6_CC_DTW_calculation” at https://github.com/JialabEleven/Vasomotion_analysis.

**Calcium and vasomotion index calculation**

For myogenic spontaneous vasomotion characterization, all data used for extracting the vasomotion index must follow a normal distribution. The vasomotion index was defined as the statistics on the total number of vasomotion events that exceeded the double standard deviation (SD) of the baseline[20]. The selection of baselines is the most critical step in analyzing calcium and vasomotion events. We determined $F_0$ by three different methods: calculating the mean value of diameter change time series curve (200-point length) as baseline which we will refer to as "$F_0$ average", calculating the minimal value of diameter change time series curve as baseline called "$F_0$ minimal" and finding the eighth smallest number in a forty sliding window producing a variable $F_0$ time series, referred to as the "$F_0$ smooth". When the amplitude of vasomotion exceeded the double SD of the baseline, it is defined as a single event. The kinetic quantification index of events including the calcium index, diameter index, and radius index. These indexes all contained parameters of frequency, SD of peak intervals (interval SD), absolute amplitude and amplitude ratio, as follows:

$$\text{Frequency:} f = \frac{n_{events}}{\mathrm{t}_{total}} \tag{12}$$

$$\text{Interval SD:} S = \sqrt{\frac{\sum_h^H t_{\Delta e}}{H-1}} \tag{13}$$

$$\text{Absolute amplitude:} \Delta D(\Delta R, \mathit{\Delta F}) = D(R, F) - D_0(R_0, F_0) \tag{14}$$

$$\text{Amplitude ratio:} \frac{\Delta D(\Delta R, \mathit{\Delta F}\ )}{D_0(R_0, F_0)}. \tag{15}$$

Among them, frequency $f$ means the all events ($n_{events}$) obtained according to the above judgment indicators divided by the total time stacks ($\mathrm{t}_{total}$, 200-point length). The interval SD ($t_{\Delta e}$ represents the time interval between each two consecutive events and H represents the amount of events interval number) shows the stability of the time interval between two consecutive events. For the calcium index, when the calcium events come, the absolute amplitude represents the changes in GCamp6s fluorescence intensity ($\Delta F$) of this time frame, and the amplitude ratio displayed as a change in calcium ($\Delta F$) over baseline ($\Delta F_0$) of this time frame. For the diameter or radius index, the average length calculated from the mean of total time length change stacks (200-point length), the absolute amplitude represents the change in width values obtained from the two boundaries ($\Delta D$ or $\Delta R$) when an event occurs, and the amplitude ratio indicates the change in width values over baseline. The inclusion criteria for the calcium events were similar as in the events selection in vasomotion. The codes can be found in “7_Vasomotion_index” at https://github.com/JialabEleven/Vasomotion_analysis.
